# The Transcriptional Logic of Mammalian Neuronal Diversity

**DOI:** 10.1101/208355

**Authors:** Ken Sugino, Erin Clark, Anton Schulmann, Yasuyuki Shima, Lihua Wang, David L. Hunt, Bryan M. Hooks, Dimitri Tränkner, Jayaram Chandrashekar, Serge Picard, Andrew Lemire, Nelson Spruston, Adam Hantman, Sacha B. Nelson

## Abstract

The mammalian nervous system is constructed of many cell types, but the principles underlying this diversity are poorly understood. To assess brain-wide transcriptional diversity, we sequenced the transcriptomes of the largest collection of genetically and anatomically identified neuronal classes. Using improved expression metrics that distinguish information content from signal-to-noise-ratio, we found that homeobox transcription factors contain the highest information about cell types and have the lowest noise. Genes that contribute the most to neuronal diversity tend to be long and enriched in factors specifically involved in neuronal function. Genome accessibility measurements reveal that long genes have more candidate regulatory elements arrayed in more distinct patterns. These elements frequently overlap interspersed repeats (mobile elements) and the pattern of repeats is predictive of gene expression. New regulatory sites resulting from elongation of neuronal genes by mobile elements may be an evolutionary force enhancing nervous system complexity.

## Introduction

The extraordinary diversity of vertebrate neurons has been appreciated since the proposal of the neuron doctrine ***(Cajal, 1888).*** Typically, this diversity is characterized by neuronal morphology, physiology, molecular expression, and circuit connectivity. The exact number of neuronal cell types remains unknown, but estimates of 40-60 have been provided for the retina ***(Macosko et al., 2015; Masland, 2004)*** and for mouse cortex (***Tasic et al., 2016; Zeisel et al., 2015).*** If similar numbers are discovered in most brain regions, the number could be in the thousands or more. Although neuronal diversity has long been recognized, the question of how this diversity arises is only beginning to be addressed ***(Arendt, 2008; Muotri and Gage, 2006).*** Describing the cell types of the brain and understanding the principles governing their diversity are fundamental goals for neuroscience.

Currently two techniques dominate the efforts to profile the transcriptional diversity of cell types in the brain: one is RNA-seq from single neurons, (single-cell RNA-seq; SCRS), (e.g. ***Shapiro et al., 2013)*** and the other is from genetically or anatomically marked pools of neurons (e.g. ***Okaty et al., 2015; Cembrowski et al., 2016).*** An obvious advantage of the SCRS approach is that, by definition, each measurement comes from only a single cell type. However, SCRS measurements can be noisy and, depending on the approach, can have limited depth and sensitivity ***(Parekh et al., 2016; Svensson et al., 2017).*** So far, the field attempts to generate accurate and precise transcriptional profiles of cell types by clustering and then averaging the profiles of single cells. But the process of clustering itself can add noise ***(Ntranos et al., 2016),*** and the unbiased nature of the measurement complicates the assessment of reproducibility. Pooling reduces noise, but can suffer from unknowingly lumping together more than one cell type. In the end, performing both methods will allow for a more confident assessment of the cell types of the brain. While large, unbiased single cell efforts have been completed or are underway, similar large scale efforts for genetically identified neurons have yet to be reported. We performed RNA-seq on the largest set to date of genetically identified and fluorescently labeled pooled neurons from micro-dissected brain regions. In total, we profiled 179 neuronal cell types and 15 non-neuronal cell types and quantitatively compared our cortical profiles to those obtained in SCRS studies. (A more precise description of our use of the term “cell type” is provided in the Methods). The comparison reveals a comparable level of homogeneity, but a much lower level of noise in the bulk sorted profiles. We have curated these reproducible and precise expression profiles to serve as a look-up table for linking single cell and cell type expression profiles to genetic strains in which they can be repeatedly accessed.

Cell types are typically identified by performing differential expression analyses. Standard differential expression methods focus on signal variance but are influenced by both information content and robustness of differential expression. We introduced two simple metrics to separate out these features of the data. Signal contrast (SC) is a signal-to-noise ratio that (unlike ANOVA) is not sensitive to differences in information content. Differentiation index (DI) is a measure of information content closely related to mutual information. Using these metrics, we identify homeobox transcription factors (TF) as the gene family with the lowest noise and highest ability to distinguish cell types and use these and other TFs to construct a compact “code” for profiled neuronal cell types. We find that the effector genes carrying the most information about cell types are synaptic genes like receptors, ion channels and cell adhesion molecules. Interestingly, a common feature of these genes is their long genomic length, reflecting the increased number and length of their introns. Our ATAC-seq results indicate that long genes contain a larger number of candidate regulatory regions which are arrayed in more diverse patterns than found in short genes, suggesting the longer length of the genes may permit increased regulatory complexity. Moreover, these long genes are elongated during evolution by insertions of mobile elements and a large portion of the candidate regulatory regions identified by ATAC-seq overlap with these mobile elements. Thus, the increased length of neuronal genes may provide a platform for evolution to fine-tune gene expression and thus diversify the cell types of the nervous system.

## Results

### A dataset of cell type-specific neuronal transcriptomes

To begin exploring the diversity in the nervous system, we collected transcriptomes from 166 types of neurons and 15 types of non-neuronal genetically/retrogradely labeled cell populations (Table 1; Figure 1 Supplement 1; Supplementary Table 1,2). Data from 9 previously published hippocampal cell types ***(Cembrowski et al., 2016),*** 2 hypothalamic cell types ***(Henry et al., 2015),*** and 2 neocortical cell types ***(Shima et al., 2016),*** harvested and processed in the same way as other samples, were also included in our analyses. Each neuron type collected represents a group of fluorescently labeled cells dissociated and sorted from a specific micro-dissected region of the mouse brain or other tissue. In most cases, the fluorescent label was genetically expressed in a mouse driver line, but retrograde labeling was used in some cases. The pipeline for cell type-specific transcriptome collection is depicted in Figure 1A (see Methods for additional details). Mouse lines were first characterized by generating a high resolution atlas of reporter expression (Figure 1B), then regions containing labeled cells with uniform morphology were chosen for sorting and RNA-seq. This effort constitutes the largest and most diverse single collection of genetically identified cell types profiled by RNA-seq. The processed data, including anatomical atlases, RNASeq coverage, and TPM are available at http://neuroseq.janelia.org (Figure 1C).

**Table 1.**
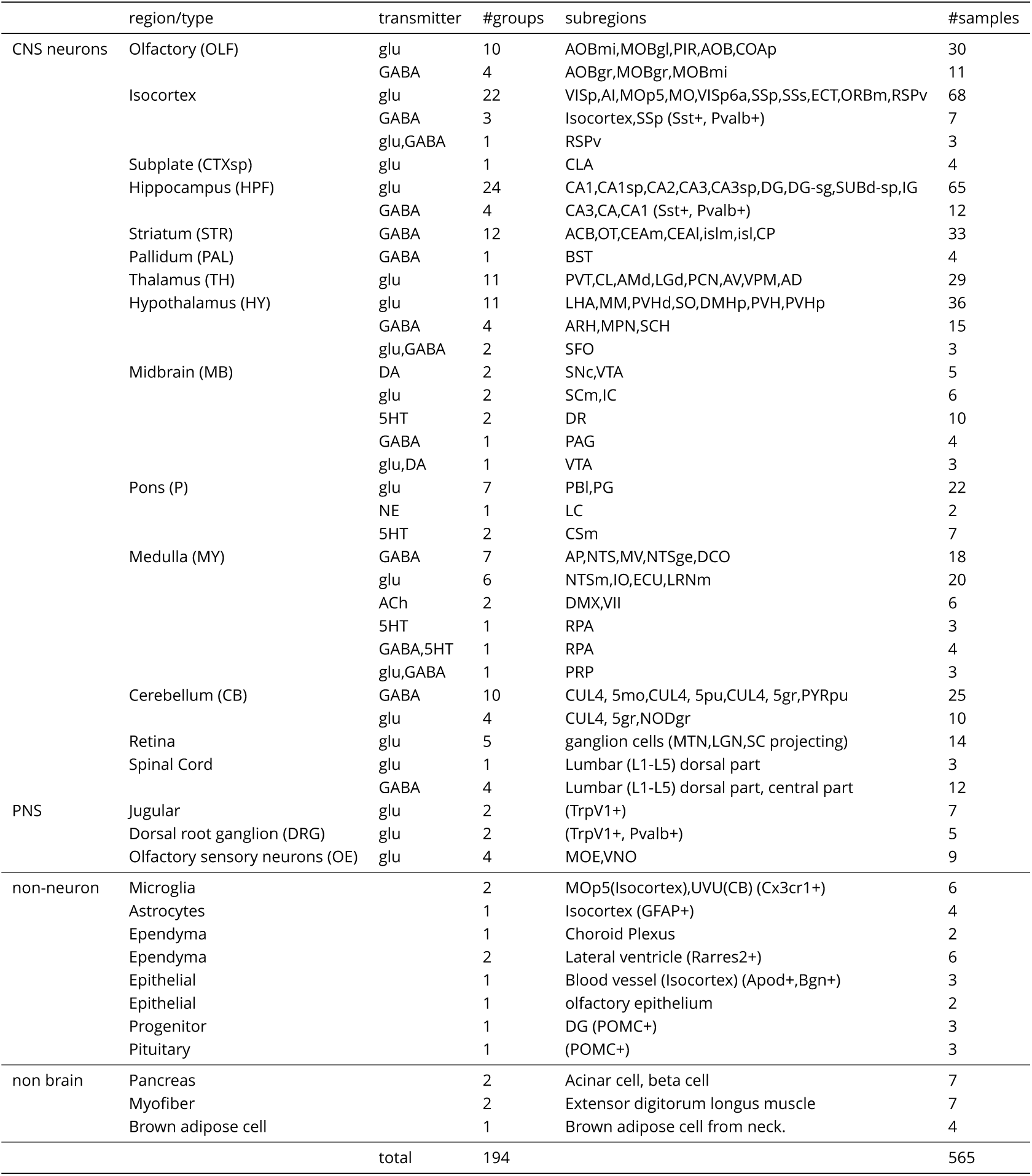
Summary of Profiled Samples.

**Figure 1.**
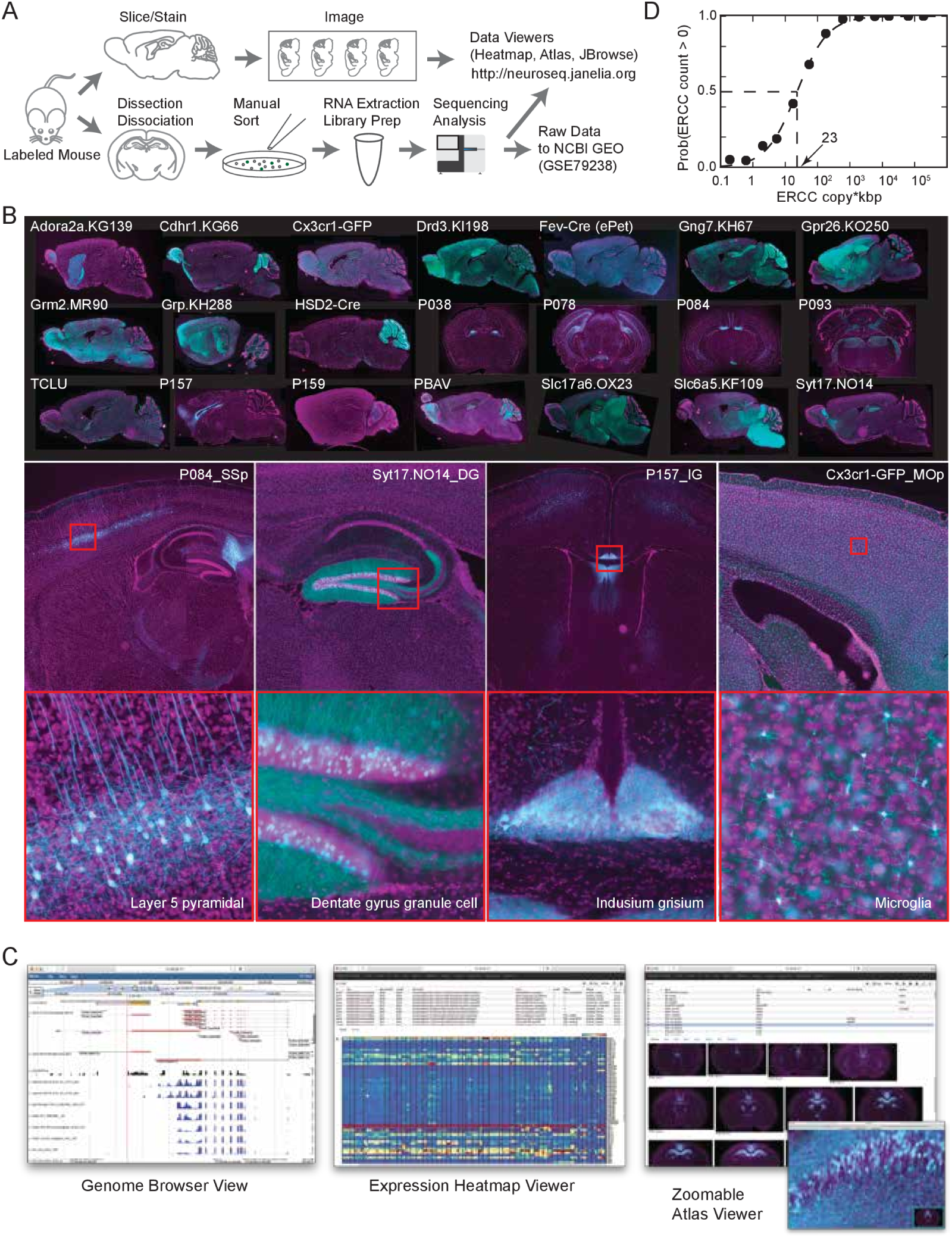
The NeuroSeq dataset. (A) Schema of pipeline for anatomical and genomic data collection. **(B)** Example sections from atlases at low (top), medium (middle) and high (bottom) magnif**i**cations. **(C)** Web tools available at http://neuroseq.janelia.org

To determine the sensitivity of our transcriptional profiling, we used ERCC spike-ins. Amplified RNA libraries had an average sensitivity (50% detection) of 23 copy*kbp of ERCC spike-ins across all libraries (Figure 1D). Since manually sorted samples had 132±16 cells (mean± sem, all following as well), this indicates our pipeline had the sensitivity to detect a single copy of a transcript per cell 80% of the time. In total we sequenced 2.3 trillion bp in 565 libraries. Total reads per library was 41 ±0.5M reads (Figure 1 Supplement 2A top). Using the aligner STAR (Dobin et al., 2012), 68.9±0.37% of the reads mapped uniquely to the mm10 genome, 2.8±0.06% mapped to multiple loci, 5.6±0.14% did not map to mm10, and 22.7±0.36% contained abundant sequences such as ribosomal RNA or mitochondrial sequences (Figure 1 Supplement 2A bottom) and 0.06%± 0.004% contained short reads (less than 30bp after removing adaptor sequences). Sequenced library data were deposited in NCBI GEO (accession number:GSE79238). This high sensitivity allowed for deep transcriptional profiling in our diverse set of cell types.

To assess the extent of contamination in the dataset, we checked expression levels of marker genes for several non-neuronal cell types (Figure 1 Supplement 2B). As previously shown ***(Okaty et al., 2011),*** manual sorting produced, in general, extremely clean data.

To demonstrate the utility of the dataset, made possible by its broad sampling of cell types, we extracted pan-neuronal genes (genes expressed commonly in all neuronal cell types but expressed at lower levels or not at all in non-neuronal cell types; Figure 1 Supplement 3). Broad sampling is essential to avoid false positives ***(Zhang et al., 2014b; Mo et al., 2015; Stefanakis et al., 2015).*** Extracted pan-neuronal genes contain well known genes such as ***Eno2*** (Enolase2), which is the neuronal form of Enolase required for the Krebs cycle, ***Slc2a3*** (chloride transporter) required for inhibitory transmission, and ***Atp1a3*** (ATPase Na+/K+ transporting subunit alpha 3) which belongs to the complex responsible for maintaining electrochemical gradients across the membrane, as well as genes not previously known to be pan-neuronal, such as ***2900011O08Rik*** (now called Migration Inhibitory Protein;***Zhang et al. (2014a)).*** Synaptic genes are often differentially expressed among neurons, but some included in this pan-neuronal list such as ***Syn1, Stx1b, Stxbp1, Sv2a,*** and ***Vamp2*** appear to be common components required in all neurons, highlighting essential parts of these complexes. Thus, this pan-neuronal gene list reveals components necessary for any neuron. The dataset should also be useful for many other applications, especially those requiring comparisons across a wide variety of neuronal cell types.

### Comparison to single cell datasets

Pools of sorted neurons may be heterogeneous if multiple neuronal subtypes are labeled in the same brain region of the same strain. SCRS has recently emerged as a viable method for profiling cellular diversity that does not suffer from this limitation. However, since profiles of cell types in SCRS studies are obtained by clustering individual, often noisy, cellular profiles, inaccuracies can arise from misclustering or overclustering. In order to assess the relative cellular homogeneity of our sorted samples, we compared the current dataset to the cluster profiles from SCRS studies. We focused on neuronal and non-neuronal cell types in the neocortex, profiled in two recent studies ***(Tasic et al., 2016; Zeisel et al., 2015).*** Assuming each sorted population corresponds to a linear combination of one or more SCRS profiles, we assessed homogeneity by linear decomposition using non-negative least squares (NNLS). We performed multiple checks on the validity of the procedure (see Figure 2 Supplements 1-3 and Methods) and found that it is able to fairly accurately decompose mixtures of component expression profiles when those components are well separated.

For each sorted cell type, the procedure identifies the weights (coefficients) of component clusters (cell types) from the SCRS datasets (Figure 2A). As expected, cell types present in the SCRS studies, but not profiled in NeuroSeq, (e.g. L4 neurons, VIP interneurons and oligodendrocytes), were not matched (purely blue columns in Figure 2A). Other cell types matched perfectly to a single SCRS cell type (e.g., microglia, astrocytes, ependyma) or matched to more than one, implying heterogeneity in the sorted profiles or poor separation of the SCRS profiles. Profiles with imperfect matches usually matched closely related cell types. For example, the NeuroSeq Pvalb interneuron group matched one or two of the SCRS Pvalb-positive interneuron clusters, and layer 2/3 (L2/3) pyramidal neurons matched SCRS L2/3 clusters, or an adjacent cluster in L4 (Tasic: L4 Arf5). The spread of coefficients repeatedly involved the same few SCRS cell clusters (e.g. columns L5b Tph2 and L5b Cdh13 in Tasic; and S1PyrL5, S1PyrL6 in Zeisel), which could occur if these clusters are not well separated, which we confirmed by a cross-validation procedure (Figure 2 Supplement 3). We measured the “purity” of the decomposition as the fractional match to the highest coefficient. The purity scores for the decomposition of NeuroSeq cell types by the two SCRS datasets were higher than those obtained for SCRS cell clusters decomposed by the other SCRS data set (Figure 2B,C). This implies that although sorted sample heterogeneity may exist in some of our sorted samples, it is comparable (or smaller) than the inaccuracies introduced by clustering single cell profiles. We also compared the separability of cell types assayed in the sorted and SCRS datasets (Figure 2 Supplement 4) by calculating the gene expression distances between each cell type within each dataset. NeuroSeq profiles were far more separable than clusters in either SCRS dataset, likely because of the noise reduction achieved by averaging across cells and because of the larger numbers of cells and reads comprising each profile. Hence sorted and single cell techniques have complimentary strengths and cross referencing both data modalities may provide the most accurate assessment of cell type specific expression.

**Figure 2.**
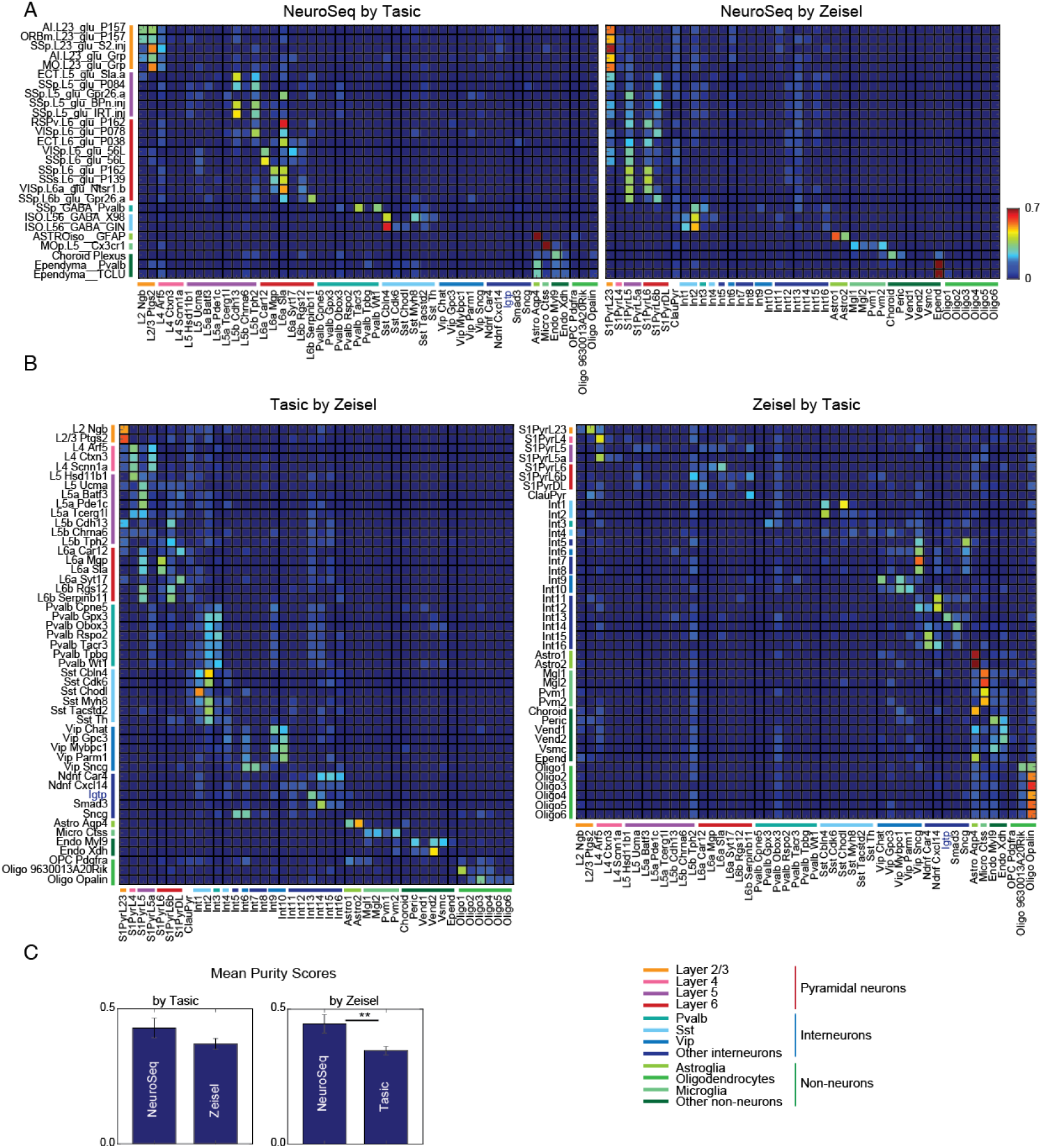
Decomposition by NNLS. (A) NNLS coefficients of NeuroSeq cell types by two SCRS datasets. **(B)** (Left) Tasic et al. clusters decomposed by Zeisel et al. clusters. (Right) Zeisel et al. clusters decomposed by Tasic et al. clusters. There are few perfect matches. **(C)** Mean purity scores for NeuroSeq and SCRS datasets. The purity score for a sample is defined as the ratio of the highest coefficient to the sum of all coefficients. (**:p<0.01, t-test.)

### Improved metrics to quantify differential expression

Analysis of expression differences between individual groups is the basis of most profiling efforts. Variance-based metrics, such as Analysis of Variance (ANOVA) F-Value or coefficient of variation (CV) are commonly used for this purpose. These metrics are jointly affected by the information content of the differential expression (pattern) and the robustness of the differences (effect size) and so cannot readily separate these two parameters. As a complement to traditional metrics and to begin mining our extensive and complex dataset for novel insights, we developed two easily calculated metrics that better separate the information content and the robustness of expression differences.

First, in order to extract the transcriptional signals related to cell type identity, we quantified each gene's ability to differentiate each pair of profiled cell types. Based on expression levels and variability (Figure 3A; Methods) we compiled a Differentiation Matrix (DM) with elements equal to one or zero depending on whether or not the gene is differentially expressed between each pair of profiles (see Methods). The Differentiation Index (DI) is simply the fraction of pairs distinguished, excluding self-comparisons; and ranges from 0 to 1. The maximum observed value of 0.65 indicates that the gene distinguishes 65% of the pairs, while a value of 0 indicates that the gene distinguishes none (i.e., expressed at similar levels in all cell types).

**Figure 3.**
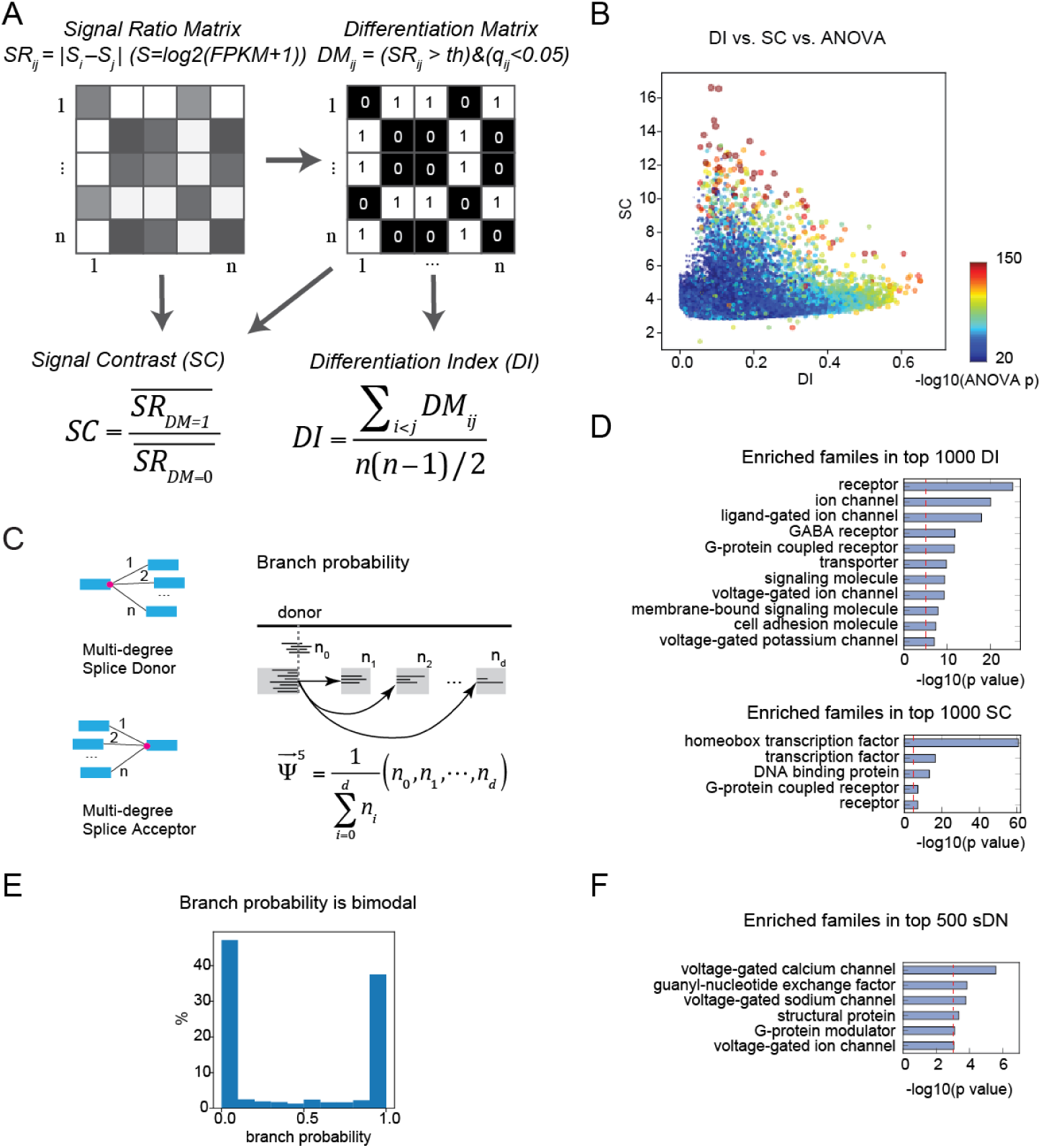
Gene expression metrics related to information content and robustness. **(A)** Expression differences between cell types are compiled into a signal ratio matrix (SR) and binarized into a differentiation matrix (DM) reflecting whether each pair of cell types is distinguished (1) or not (0). The Differentiation Index (DI) is the fraction of nonzero values. The Signal Contrast (SC) is the average expression difference between distinguished pairs divided by the average expression difference between undistinguished pairs. **(B)** Highly significant ANOVA genes (warm colored dots) include a mixture of genes with high SC and low DI and genes with low SC and high DI. **(C)** Definition of generalized PSI (percent spliced in). For a splice donor, a generalized form of PSI (donor branch probability) can be defined as the joint distribution of transition probabilities from the donor to each acceptor. Acceptor branch probability can be defined conversely. **(D)** PANTHER *(Thomas, 2003)* gene families enriched in the top 1000 DI and the top 1000 SC genes. Red lines indicate the *p =* 10^-5^ threshold used to judge significance. **(E)** Histogram of all donor branch probabilities from alternatively spliced sites. The distribution is highly bimodal, indicating that alternative splicing is “all or none” for each site in each cell type (though often varying between cell types). **(F)** PANTHER gene families enriched in the top 500 DN genes. The number of cell types distinguished by a gene's splice variants (sDN; see Methods for calculation) rather than the ratio (DI) is used since the denominator of DI (total number of cell types potentially distinguished) varies for each gene. This is because genes not expressed in a cell type can contribute to distinctions based on expression, but not to those based on splicing. Red lines indicate the *p =* 10^-3^ threshold used to judge significance.

The ability to detect transcriptional differences between cell types depends on both magnitude of difference and associated noise. To quantify this in our second metric, we defined the Signal Contrast (SC), which closely reflects Signal-to-Noise-Ratio (SNR). Since the signals we are interested in are the gene expression differences distinguishing cell types, we used a noise estimate derived from all undistinguished pairs from the same gene. SC, which indicates how robustly pairs are distinguished, is the ratio of the average effect size for distinguished and undistinguished pairs. High SC genes robustly distinguish cell populations and are therefore suitable as "marker genes".

Our metrics outperform existing metrics such as ANOVA, CV, and Fano factor in distinguishing the information content and robustness of differential expression. To illustrate the properties of DI and SC relative to existing metrics, we calculated these metrics against various simulated expression patterns with added noise (Figure 3 Supplement 1A). The results (Figure 3 Supplement 1A, lower part) demonstrate that DI (blue) is highly correlated with mutual information (MI; green), yet much easier to calculate. This makes intuitive sense, since the division of cell types into those that can and cannot be distinguished (DM; Figure 3A) corresponds to a unit of information about cell types provided by a gene expression pattern (for more details of the relationship between DI and MI, see Figure 3 Supplement 1C and 2). The simulations also show that DI is fairly independent from SNR. For example, both high and low SNR binary patterns yield similar DIs. In contrast, SC (orange) is independent from MI, but is highly correlated to SNR. Thus, DI provides an estimate of the information content of expression patterns across cell types, whereas SC provides an estimate of SNR.

Unlike DI and SC, traditional variance-based methods like ANOVA F-values and CV are either affected by both MI and SNR (ANOVA) or by neither (CV). These differences between metrics are summarized in Figure 3 Supplement 1B. The fact that ANOVA does not distinguish between information content and SNR is also apparent in the data. As shown in Figure 3B, high-ANOVA genes include both high DI and high SC genes. Therefore, SC and DI are useful because they provide independent measures of the robustness and magnitude of differential expression between cell types.

### Genes with the highest information regarding cell types

To determine the types of genes most differentially expressed (highest DI) and most robustly different (highest SC) between cell types, we used the PANTHER ***(Thomas, 2003)*** gene families (Figure 3D). As expected, high DI genes are enriched for neuronal effector genes including receptors, ion channels and cell adhesion molecules (Figure 3D top). The highest signal-to-noise expression differences (highest SC) were those of homeobox transcription factors (TFs) and the more inclusive categories (TFs, DNA binding proteins) that encompass them (Figure 3D bottom). Hence DI and SC respectively emphasize the information content of genes mediating the distinctive neuronal phenotypes that distinguish cell types, and the robust, low-noise expression of genes involved in shaping these cell types unique transcriptional programs.

Genes may also contribute to cell type differences through differential splicing. We analyzed splicing events by computing the relative likelihood (branch probabilities) of each donor site in a transcript being spliced to multiple acceptor sites, and of each acceptor site being spliced to multiple donors (Figure 3C). Interestingly, when these branch probabilities are computed separately for each cell type, they are highly bimodal, reflecting virtually all-or-none splicing at each alternatively spliced site. This pattern has previously been observed for individual cells in some systems ***(Shalek et al., 2013).*** The present observations suggest that these splicing decisions are made at the level of cell types, rather than independently for individual cells of the same type. We applied a variant of the DM/DI method to alternative splicing (Figure 3C,E,F; for details see Methods) and found that voltage-gated calcium and sodium channels are highly alternatively spliced, consistent with previously known results (e.g. ***Lipscombe et al., 2013).*** We also found that G-protein modulators, especially guanyl-nucleotide exchange factors (GEFs), are highly alternatively spliced. Hence, differential splicing of multi-exon genes also contributes to transcriptome diversity across neuronal cell types.

SC, like SNR, is a ratio between signal and noise, and so can reflect high expression levels in ON cell types (high signal), low expression levels in OFF cell types (low noise), or both. Homeobox genes are not among the most abundantly expressed genes. Their average expression levels (~30 FPKM) are significantly lower than, for example, those of neuropeptides (~90 FPKM). This suggests that the high SC of homeobox TFs depend more on low noise than on their high signal. In fact, most homeobox TFs have uniformly low expression in OFF cell types (e.g. Figure 4A). We quantified this “OFF noise” for all genes and found that homeobox genes are enriched among genes that have both low OFF noise and at least moderate ON expression levels (red dashed region in Figure 4B).

**Figure 4.**
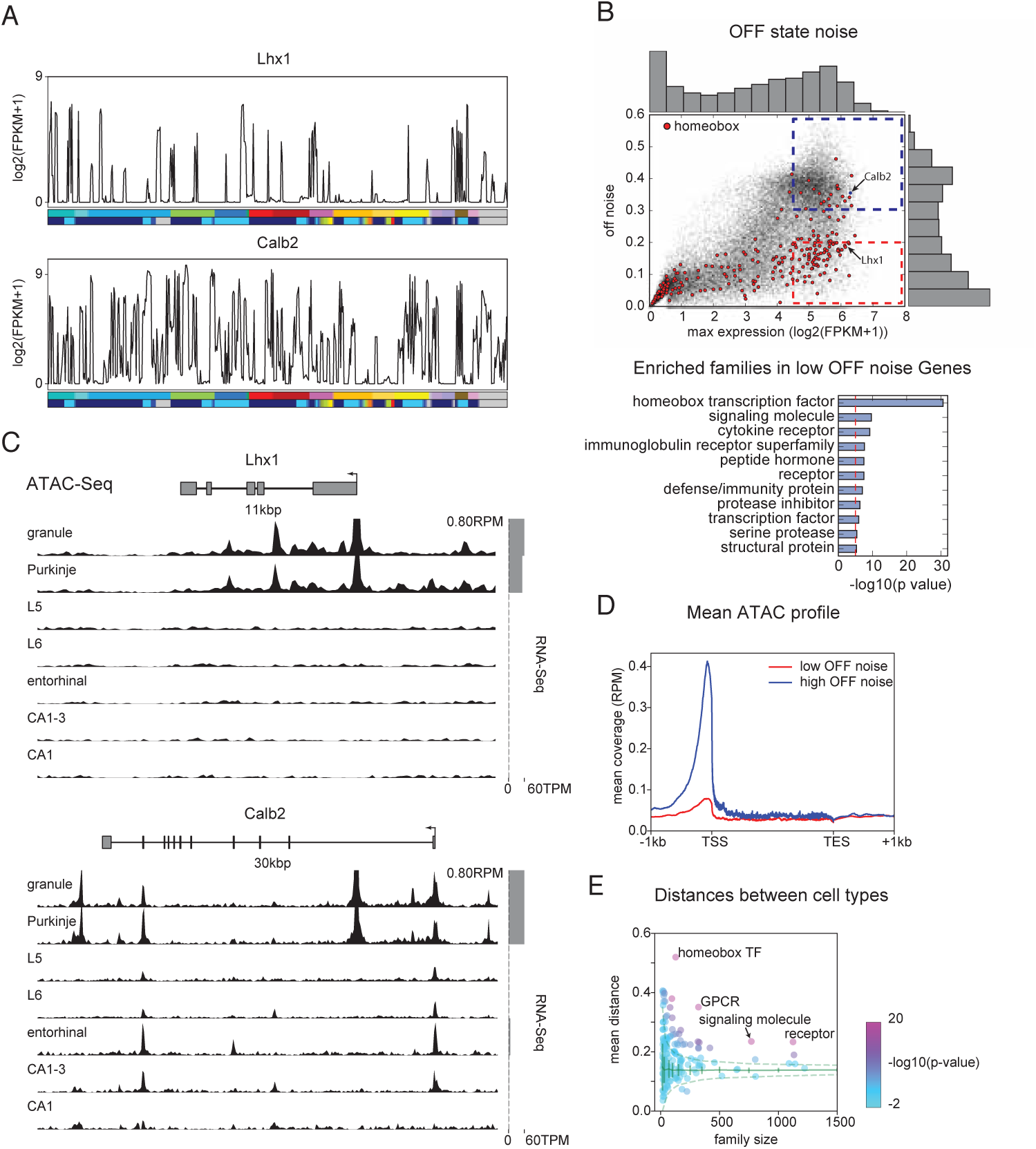
Mechanisms contributing to high information content and low noise of Homeobox TFs. (A) Example expression patterns of a LIM class homeobox TF *(Lhx1)* and a calcium binding protein *(Calb2)* with similar overall expression levels. Cell type legend is given in Figure 1 Supplement 1. **(B)** (upper) OFF state noise (defined as std. dev. of samples with FPKM<1) plotted against maximum expression. (lower) PANTHER families enriched in the region indicated by red dashed lines in the upper panel. **(C)** Average (replicate N=2) ATAC-seq profiles for the genes shown in A. Some peaks are truncated. Expression levels are plotted at right (grey bars). **(D)** Length-normalized ATAC profile for genes with high (> 0.3, blue dashed box in B, n=853) and low (< 0.2, red dashed box in B, n=1643) off state expression noise. **(E)** Mean separability of cell types for PANTHER families. Separability is a measure of gene expression distance (defined as the average of 1- Pearson's corr. coef.) calculated across a set of genes. Since dispersion of separability decreases with family size, results are compared to separability calculated from randomly sampled groups of genes (green solid lines: mean and std. dev.; green dashed lines: 99% confidence interval). Z-scores: homeobox TF: 17.4, GPCR: 16.1, receptor: 13.1 and signaling molecule: 11.2.

Since tight control of expression may reflect closed chromatin, we measured chromatin accessibility using ATAC-seq (***Buenrostro et al., 2013)*** on 7 different neuronal cell types (see Methods). As expected, compared to high-noise genes (Figure 4C bottom), genes with low OFF noise were more likely to have fewer, smaller peaks within their transcription start site (TSS) and gene body (Figure 4C top, Figure 4D), consistent with the idea that their expression is controlled at the level of chromatin accessibility.

Functionally, the tight control of homeobox TF expression levels may reflect their known importance as determinants of cell identity, and the fact that establishing and maintaining robust differences between cell types may require tight ON/OFF regulation rather than graded regulation. If they are, in fact, important “drivers” of cell type-specific differences, their expression pattern should be highly informative about cell types. However, the homeobox family was not identified on the basis of a particularly high DI (Figure 3C and Figure 4 Supplement 1B; mean DI=0.21; rank 16th) compared to, for example, cyclic nucleotide-gated ion channels (mean 0.31, highest) or GABA receptors (0.29, 2nd). We infer that this is due to the fact that graded expression differences also contribute to DI. Since binary ON/OFF expression patterns may be more critical for cell type specification than graded expression patterns, we calculated a binary version of DI (bDI; see Methods). With this metric, the homeobox TF family is the most enriched PANTHER family among the top 1000 bDI genes (Figure 4 Supplement 1A) and had the 2nd highest average bDI (0.07) among PANTHER families after neuropeptides (0.08) (Figure 4 Supplement 1B). Among TF subfamilies, the LIM domain subfamily of homeobox genes had the highest mean bDI (Figure 4 Supplement 1C), consistent with its known role in specifying spinal cord and brainstem cell types ***(Tsuchida et al., 1994; Philippidou and Dasen, 2013).***

The ability of gene families to provide information about cell types is determined by both how informative individual family members are, and the relationships between them. If the information across family members is independent, the overall information is increased relative to the case in which multiple members contain redundant information (Figure 4 Supplement 1D). This aspect of “family-wise” information is not captured by “gene-wise” metrics like mean bDI, or by enrichment analysis (Figure 3C, Figure 4 Supplement 1A-C). One way of capturing the additive, non-redundant information within a gene family is to measure its ability to separate cell types using a distance metric. This analysis (Figure 4E) reveals that homeobox TFs yield the largest distances between cell types. Thus, homeobox TFs provide the best separation of profiled cell types both individually (Figure 4 supplement 1A,B) and as a family (Figure 4E). It has long been known that a subset of homeobox TFs, the HOX genes, play an evolutionarily conserved role in specifying cell types in invertebrates ***(Kratsios et al., 2017; Zheng et al., 2015)*** and in the vertebrate spinal cord and brainstem ***(Dasen and Jessell, 2009; Philippidou and Dasen, 2013).*** Our current analyses suggest that the larger family of homeobox TFs play a broader role in transcriptional diversity of cell types across the mammalian nervous system.

In summary, by defining novel metrics DI and SC, we identify homeobox TFs as the most robustly distinguishing family of genes as well as synaptic and signaling genes as the most differentially expressed genes. These two categories of genes drive neuronal diversity by orchestrating cell type-specific patterns of transcription and by endowing neuronal cell types with specialized signaling and connectivity phenotypes.

### A compact TF code for neuronal identity

In addition to identifying the most informative transcription factors across the entire set of cell types studied, we also identify the most informative TFs for individual cell types. To accomplish this, we extracted the most compact set of “ON” or “OFF” TFs needed to specify each cell type generating a hierarchy of TFs constituting a decision tree that efficiently classifies cell types ***(Gabitto et al., 2016).*** At each level of the tree, TFs were chosen to optimally bisect (by their expression level) the set of cell types into two groups that differed maximally from each other in terms of their overall expression profile (assessed within the full transcriptome). To generate a classifier operating at each level of anatomical organization, we favored TFs whose bisected groups are consistent with anatomical divisions (see Methods for details).

The selected TFs included many genes previously implicated as key transcriptional regulators (KTRs) in the development or maintenance of the distinguished cell types. For example, *Foxg1,* which split forebrain from other cell types, is known to be critically required for normal development of the telencephalon ***(Xuan et al., 1995; Danesin and Houart, 2012)*** and is known to function cell autonomously within the olfactory placode for the production of olfactory sensory neurons, as well as for all other cells in the olfactory lineage ***(Duggan et al., 2008).*** Similarly, at the next levels, *Tbr1 **(Bedogni et al., 2010)**, Satb2 **Leone et al. (2014)**, Egr3 **(Chandra et al., 2015)**, Isl1 **(Lu et al., 2013)*** and *Emx2 **(Zhang et al., 2016)***, are known as KTRs involved in the development and/or maintenance of the relevant cell types, providing significant validation of this method.

The TF code identified for each cell type is not unique. First, there are additional TFs that are consistent with the tree (see Supplementary Table 3). Second, past the first level *(Foxg1),* TFs may be expressed outside of the cell types shown and so could contribute to encoding other expression differences. More generally, the details of the tree may depend on the precise procedure used to extract it. We explored variant procedures that better preserved the known anatomical and developmental relationships between cell types (Figure 5 Supplement 1) as well as procedures that made no assumptions about these relationship whatsoever (Figure 5 Supplement 2). Interestingly, in each case, the majority of the same genes were identified, suggesting they encode cell type information that is robust to the precise methods used to extract them.

**Figure 5.**
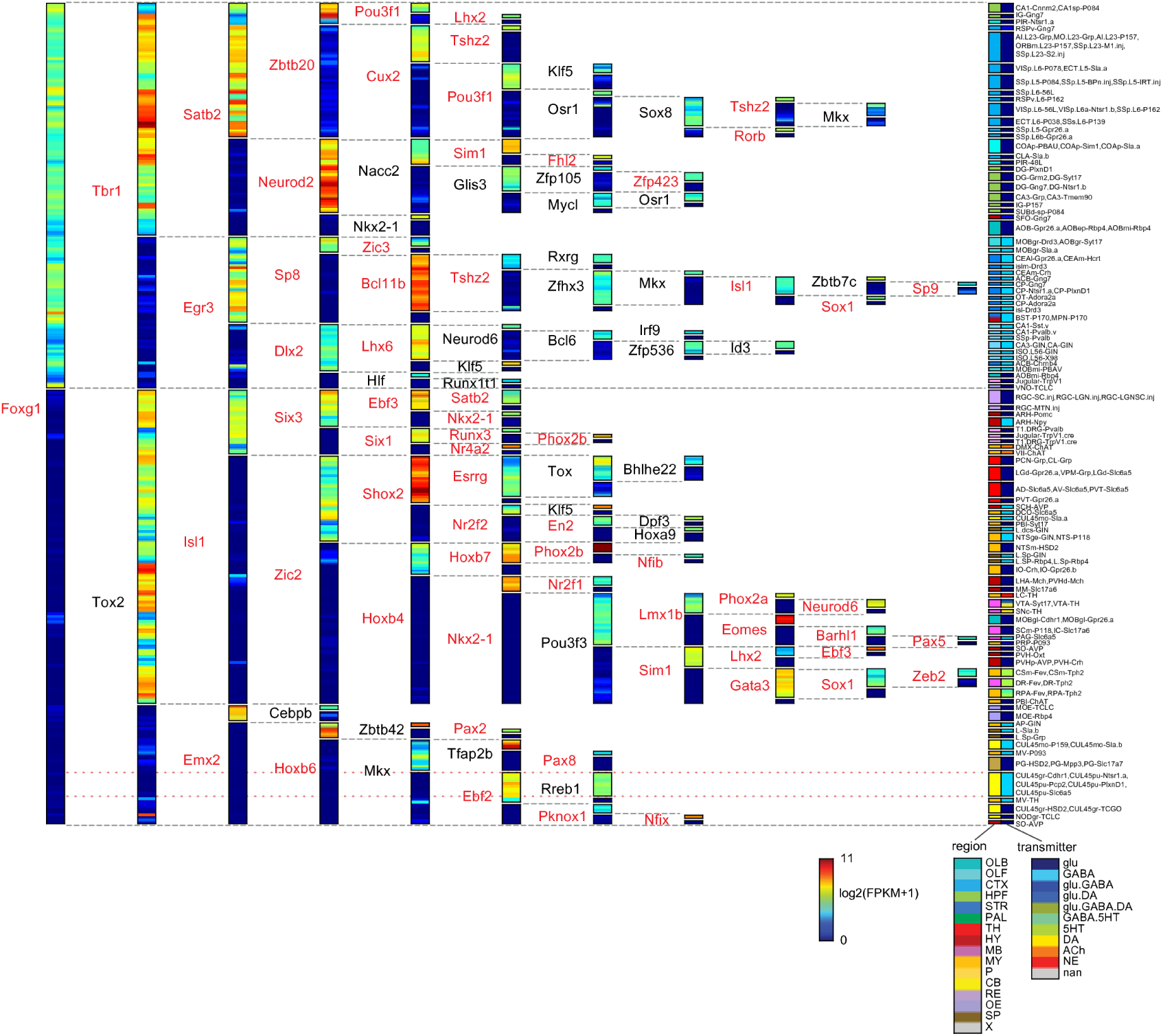
A compact TF code. A decision tree classifier constructed from the most informative TFs for profiled cell types. Cell types are bisected at each node by TF expression level, (color scale). Each cell type can be specified by the “ON” (warm colors) or “OFF” (cool colors) expression of 4 to 11 TFs as indicated. For example, Purkinje cells (yellow-light blue group near the right bottom corner, consisting of CUL4,5gr-Cdhr1, CUL4,5pu-Pcp2, etc.) have a code which can be read from left to right within the red dotted lines, consisting of: Foxg1(OFF)-Tox2(OFF)-Emx2(OFF)-Hoxb6(OFF)-Mkx(OFF)-Ebf2(ON)-Rreb1(ON). Blue dashed lines mark positions of ON/Off transitions for each TF.

Although the decision tree classifier identifies many known KTRs, it also suggests hypotheses about less studied genes. For example, ***Tox2*** has received little prior study in the CNS, although it has recently been identified and replicated as a locus of heritability for Major Depressive Disorder ***(Zeng et al., 2016).*** Based on its position in the tree, we hypothesize that ***Tox2*** is a KTR of midbrain, hypothalamic and hindbrain cell types, including dopaminergic and serotonergic cell types in these regions, although its expression in other cell types may also contribute. Hence the tree of identified TFs is a robust and rich source of novel hypotheses about transcriptional regulation in genetically identified cell types. Known and hypothesized KTRs identified by the decision tree classifier are tabulated in Supplementary Table 3.

### Long genes contribute disproportionately to neuronal diversity

We found that neuronal effector genes such as ion channels, receptors and cell adhesion molecules have the greatest ability to distinguish cell types (highest DI; Figure 3C). Previously, these categories of genes have been found to be selectively enriched in neurons and to share the physical characteristic of being long ***(Sugino et al., 2014; Gabel et al., 2015; Zylka et al., 2015)***. Consistent with this, DI is strongly biased toward long gene length (Figure 6A). Interestingly, the expression of long genes is not uniform across brain regions, but is highest in the evolutionary newer forebrain and is lower in the older brainstem and hypothalamus (Figure 6B). Non-neuronal cell types expressed only 1/2 to 1/5 as many long genes as neuronal cell types (blue bars in Figure 6B). This was true even for non-dividing cell types like myocytes and largely non-dividing tissues like the heart (separate data not shown). Hence long genes, which are preferentially expressed in neurons, also contribute most to the differential expression between neuronal cell types.

**Figure 6.**
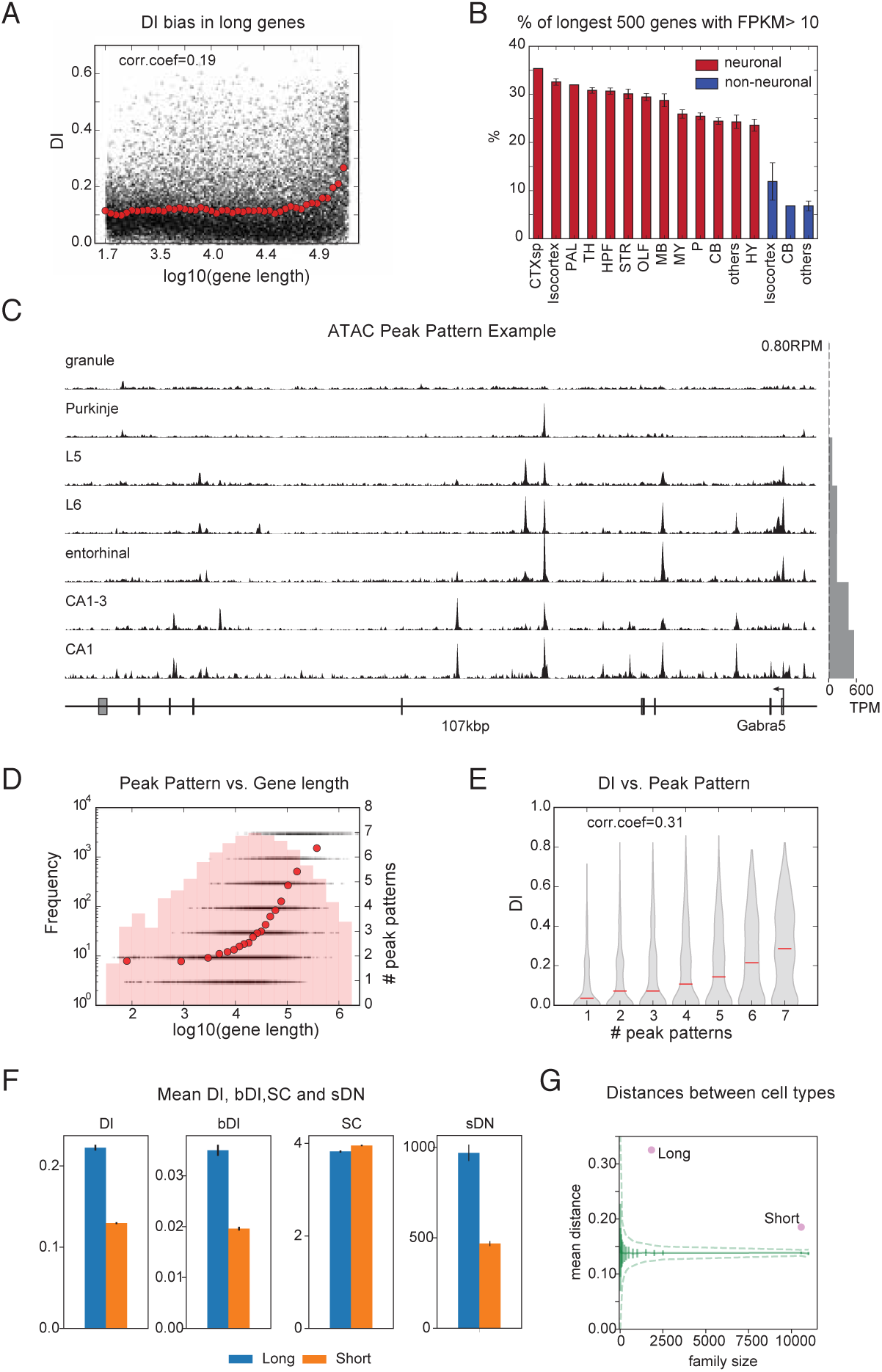
Long genes have a greater capacity for differential expression. (A) Black dots: DI of each gene is plotted against sorted gene length. Red dots: binned average of DIs (1000 genes per bin, sorted by length). **(B)** Fraction of the longest 500 genes expressed within each brain region profiled for neuronal (red bars) and non-neuronal cell types (blue bars). **(C)** ATAC-seq peaks for *Gabra5* showing different patterns of peaks for each of 7 cell types. Scale (top right) in reads per million. Expression levels for each cell type are shown at right (gray bars). **(D)** Black dots: number of distinct peak patterns observed across 7 ATAC-seq profiled cell types plotted against the gene length for each gene; 7 corresponds to a distinct pattern for each profiled cell type. Red dots: binned averages of black dots as in panel A. Background histograms show numbers of genes in each length bin. **(E)** Violin plot showing the relationship between DI and the number of different patterns of ATAC-seq peaks. Corr.coef. (0.31) is greater than that between DI and gene length (0.19; panel A). **(F)** Average metrics for long (≥100kbp) and short (<100kbp) neuronal genes (reproducibly expressed in neuronal cell types). **(G)** Separability of cell types calculated as in Figure 4E, but using long neuronal genes and short neuronal genes rather than functionally defined gene families. Z-score is 33.2 for long and 22.1 for short neuronal genes. Both are highly different from randomly sampled genes (green solid lines mean and Std. dev.; dashed lines = 99% confidence interval), but long genes provide greater separation.

REST is an important zinc-finger transcription factor that represses expression of neuronal genes in non-neurons ***(Chong et al., 1995; Schoenherr and Anderson, 1995).*** We wondered if REST preferentially targets long genes. To assess the magnitude of this effect and its influence on the length distribution of neuronal genes (Figure 6 Supplement 1A), we plotted the length-dependence of genes containing RE1/NRSE elements (Figure 6 Supplement 1B) and observed that they are indeed biased toward long genes. When these REST targets are removed from neuronally expressed genes, the length distribution of expressed genes looks similar to that of non-neurons (Figure 6 Supplement 1C). However, consistent with the fact that only 8.6% of neuronally expressed genes are REST targets (contain an NRSE), the removal of these genes has only a modest effect on the length distribution of DI (Figure 6 Supplement 1D). Therefore, although REST targets are long, many other long genes also contribute to neuronal diversity.

Long genes differ from more compact genes primarily in the number and length of their introns, which, for the longest genes, comprise all but a few percent of their length (Figure 6 Supplement 1E). Introns often contain ***cis*** regulatory elements that regulate transcription, splicing and other aspects of gene expression. Could these longer introns increase the regulatory capacity of long genes? In order to determine whether or not the introns of long genes have enhanced regulatory capacity, we identified candidate regulatory elements as sites of enhanced genome accessibility using our ATAC-seq data. As expected, long genes had more candidate regulatory elements (ATAC peaks; Figure 6 Supplement 1F) and these peaks were present in a greater number of distinct patterns per gene across cell types (Figure 6C,D). Consistent with the hypothesized role in differential expression,the number of unique patterns correlated well with the degree of differential expression across cell types (Figure 6E). Hence long genes have enhanced regulatory capacity that correlates with their enhanced contribution to neuronal diversity.

To compare candidate regulatory elements in long genes between neurons and non-neurons, we used publicly available DNase-seq data from the ENCODE project ***(Dunham et al., 2012).*** We found a significantly higher number of open chromatin sites in brain compared to non-brain tissue. This bias was particularly pronounced in forebrain, and was stronger in human than in mouse tissue (Figure 6 Supplement 1G-J). Together, these data support the hypothesis that neuronal genes may have increased in length over evolutionary time in part to support more complex and nuanced regulatory regimes.

To assess the relative contribution of long (≥100kbp) and short (<100kbp) genes, we first calculated averages of “gene-wise” metrics (Figure 6F). Signal contrast is comparable between these two groups of genes, but, for all other metrics (DI, bDI, sDN; for sDN see Figure 3 D,E and Methods), averages for long genes are about twice that of short genes. Enhanced alternative splicing of long genes (high sDN) is readily understandable from the increased number of alternative splice sites in long genes (Figure 6 Supplement 1K). Although there is no significant difference in SC between long and short genes, low OFF noise genes (Figure 4B-D) are significantly shorter than high OFF noise genes (Figure 6 Supplement 1L).

To assess the “group-wise” contribution (akin to the “family-wise” analysis of Figures 3C,F and 4E), we first observed that both groups are fairly decorrelated between member genes (Figure 6 Supplement 1M). Despite similar decorrelation, the distances between cell types based on long gene expression are larger than those obtained from expression of short genes (Figure 6G). Thus, long genes, as a group, contribute more than short genes to neuronal diversity.

### TE insertions elongate genes and carry regulatory information

The above results indicate that gene length is an important contributor to gene expression diversity across cell types. Gene lengths differ widely across species (Figure 7A and Figure 7 Supplement1A), suggesting genes are elongated during evolution. In fact, evolutionary older genes are longer (Figure 7 Supplement 1B; **Grishkevich and Yanai (2014)).** To better understand mechanisms of gene elongation over mammalian evolution, we examined segments inserted into the human and mouse genomes by comparing them to closely related species (Figure 7B). Plotted in Figure 7B (left) is a histogram of the lengths of the segments inserted into human (see also **Mikkelsen et al. (2005).** Two clear peaks are recognizable, corresponding to Alu and L1 repeats. Moreover, around 92% of the base pairs of the inserted segments overlap with known repeats (Figure 7B inset; **Bao et al. (2015)).** Similar results are observed in the mouse genome (Figure 7B right; see also ***Pozzoli et al. (2007)***). These comparisons indicate that genes are elongated by transposable element (TE) insertions.

**Figure 7.**
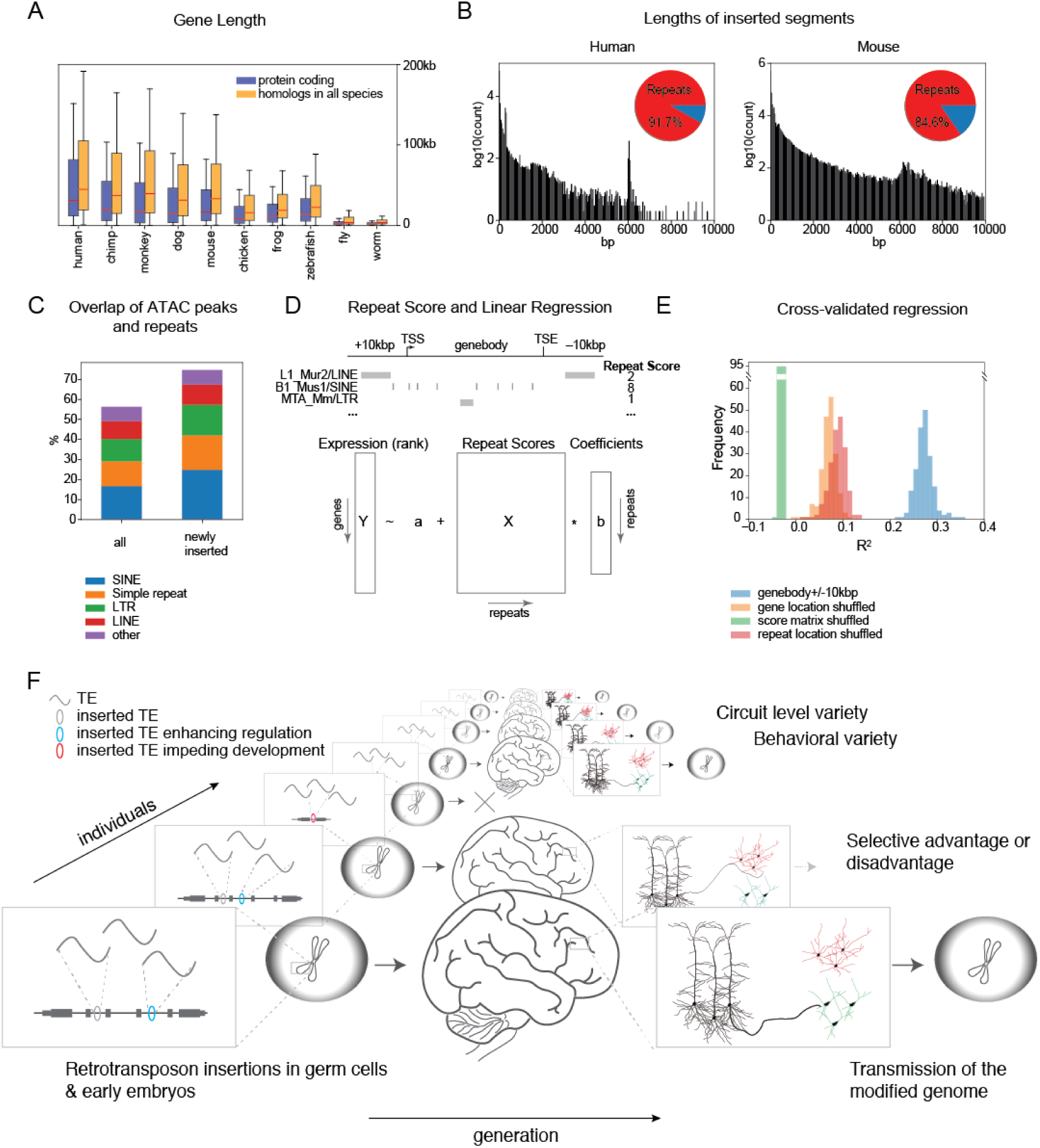
Genes are elongated by TE insertions and TEs contain information for gene expression. **(A)** Distribution of gene length for various well annotated species. Red lines indicate means and whiskers indicate inter-quartile range. Blue bars are all protein coding genes and yellow bars are the subset of genes with homologs in all species. (human: *Homo sapiens*; chimp: *Pan troglodytes*; monkey: *Macaca mulatta,* mouse: *Mus musculus*; dog: *Canis lupus familiaris;* chicken: *Gallus gallus;* frog: *Xenopus tropicalis;* zebrafish: *Danio rerio;* fly: *Drosophila melanogaster;* worm: *Caenorhabditis elegans)* **(B)** Histograms of lengths of segments inserted into the human genome compared to chimp (left) and mouse genome compared to rat (right). Peak near 300bp (more visible in human) corresponds to Alu, and near 6000bp corresponds to LINE. Pie charts (insets) indicate fraction of inserted bp overlapping transposable elements (TE) and other types of repeats. Gorilla and Guinea pig are used as surrogates of common ancestors of human and chimp, and mouse and rat, respectively (see Methods). **(C)** Percentage of ATAC peaks overlapping major categories of repeat elements. Left side: all ATAC peaks, right side: ATAC peaks overlapping recently inserted segments calculated in (B). **(D)** Schema describing repeat score and regression model. Repeat scores (upper panel) are calculated separately for each type of repeat element and for each gene as the count of that element in the specified interval determined by the gene. Regressions (lower panel) are calculated separately for each cell type by fitting coefficients (b) to ranked expression levels (Y) using intercept(a) and repeat score (X). **(E)** Fits to 80% of the genes are cross validated using the remaining 20%. Histograms show cross validated *R*^2^ for each cell type (blue), and for controls shuffling the relationship between repeat scores and genes(score matrix; green) or changing the repeat score by randomly changing the location of repeats (red) or by calculating the repeat score over a randomly selected genomic interval of the same length as the gene (orange). The latter two shuffling methods retain some predictive value compared to shuffling the repeat score matrix (green) since they maintain the correlation between gene length and expression (See Figure 7 Supplement 1C). **(F)** A model of how neuronal genes become elongated over evolutionary time scales.

Since long genes have a greater number of candidate regulatory elements, as indicated by more ATAC-peaks, we asked whether these can originate from mobile elements. As shown in Figure 7C, 56% of the ATAC peaks overlap known repeats and this number increases to 75% when only newly inserted segments are considered, indicating that TEs may carry regulatory functions. To explore the possibility that TE/repeats contribute to global regulation of neuronal gene expression, we fit gene expression levels with counts of individual repeats within and surrounding each gene (Figure 7D). The ***R^2^*** values for each cell type calculated using test genes (20%) not used for fitting (Figure 7E, blue) are much larger than expected by chance (Figure 7E, green/red/orange). If counts and genes are shuffled (green) cross validated *R*^2^ values drop below 0. However, if the length of the gene is retained in the shuffling control (orange, red) the *R*^2^ values drop to about 1/3 of those in the original fitting. This reflects the fact that gene length is highly correlated with expression (Figure 7 Suplement 1C; c=0.418: mean Pearson’s r between log gene length and expression rank) and some repeats, such as SINEs, are highly correlated with both gene length (c=0.841) and expression (Figure 7 Supplement 1C; mean c=0.454). We also varied the size and position of the regions used to count repeats and found that predictions about expression (***R***^2^) were best when including the gene body and the adjacent 10 ~ 50kbp. (Figure7 Supplement 1D,E). There are several prior examples relating TE regulation of gene expression (e.g. ***Han et al. (2004)***; **Chuong et al. (2016)**), however, this is the first to show the existence of a global network of TEs affecting gene expression in neurons.

In summary, genes are elongated by insertions of TEs which overlap candidate regulatory elements, and are predictive of relative gene expression levels, suggesting they may increase the capacity of long genes to be differentially expressed.

## Discussion

### A Resource of Neuronal Cell type specific Transcriptomes

The dataset presented here is the largest collection of cell type-specific neuronal transcriptomes obtained by RNA-seq (Table 1) and so offers the broadest view to date of the transcriptional basis of neuronal diversity. Prior RNA-seq data from sorted cells have been focused primarily on what distinguishes neurons as a class from other brain cell types ***(Zhang et al., 2014b)***, or have focused on a limited number of brain regions, such as the somatosensory cortex, hippocampus ***(Zeisel et al., 2015; Cembrowski et al., 2016; Tasic et al., 2016)*** and retina ***(Macosko et al., 2015).*** Our strategy of profiling labeled populations of ~ 100 cells is intermediate between single cell profiling, which can be limited by the noisiness of single cell assays ***(Marinov et al., 2013)*** and tissue profiling, which cannot resolve the heterogeneity of component cell types ***(Nelson et al., 2006).*** This approach enabled us to obtain highly sensitive and reproducible transcriptomes from genetically accessible target populations. The wide range of cell types in the dataset is suitable for addressing general questions regarding neuronal identity and diversity, but at the same time, the fact that each transcriptome corresponds to a genetically (or retrogradely) labeled population, allows investigation of the same population of the cells across time and labs in order to address more specific questions about those cell types.

We developed a quantitative approach for comparing cell type profiles across multiple studies using NNLS decomposition. The results reveal multiple cases in which pooled cell profiles mapped to more than one SCRS profile. It is likely that at least some of these cases represent biologically distinct cell types that share a genetic marker (like subtypes of Pvalb interneurons). However, in most of these cases, the SCRS clusters were barely separable, and the two SCRS studies available for comparison did not agree. Given the complimentary advantages of improved reproducibility, separability and deeper depth of sequencing afforded by the pooling approach, and of reduced heterogeneity afforded by the SCRS approach, it is likely that further integration of these approaches with other modalities, such as FISH ***(Moffitt et al., 2016)*** will be needed to accurately catalog the full census of brain cell types.

### A transcriptional code for neuronal diversity

We developed novel, easily calculated metrics that capture essential features of the robustness and information content of transcriptome diversity. These measures are not cleanly captured by traditional variance-based metrics like ANOVA and CV (Figure 3 Supplement 1). We found that the homeobox family of TFs exhibited the most robust (high SC) expression differences across cell types (Figure 3D bottom). These ON/OFF differences were characterized by extremely low expression in the OFF state (Figure 4A-D). Mechanistically, the low expression was associated with reduced genome accessibility measured by ATAC-seq (Figure 4C,D), presumably reflecting epigenetic regulation, known to occur for example at the clustered Hox genes via Polycomb group (PcG) proteins ***(Montavon and Soshnikova, 2014).*** Although this regulation has been studied most extensively at Hox genes, genome-wide ChIP studies reveal that PcG proteins are bound to over 100 homeobox TFs in ES cells ***(Boyer et al., 2006).*** Our results indicate that strong cell type-specific repression persists in the adult brain. Presumably this represents the continued functional importance of preventing even partial activation of inappropriate programs of neuronal identity.

As a group, homeobox TFs distinguished 98% of neuronal cell types profiled. Historically, homebox TFs are well known to combinatorially regulate neuronal identity in Drosophila and C. elegans (***Kratsios et al., 2017)*** and the vertebrate brainstem and spinal cord (***Dasen and Jessell, 2009; Philippidou and Dasen, 2013).*** The continued expression of homeobox TFs throughout the adult mammalian nervous system suggests that they likely also contribute to the maintenance of neuronal identity.

In order to reveal the relationship between specific cell types and TFs, we constructed a TF decision tree for classifying profiled cell types. As expected from their high information content, homeobox TFs figured prominently in this list (49/127). Many of the identified factors are known to be key transcriptional regulators of the cell types in which they continue to be expressed (Supplemental Table 3). In most cases it is not known whether or not these roles occur only in development, or are also important for the maintenance of neuronal identity. Lists of expressed TFs and the genetically accessible cell types in which they are expressed provide a ready source of testable hypotheses about how cell type specific transcriptional identity is maintained in the adult nervous system.

### Long genes shape neuronal diversity

Our study suggests that long genes contribute disproportionately to neuronal diversity (Figure 6A,F,G). Increases in the number of alternative start and splice sites present in longer genes increase neuronal diversity (Figure 6F), but in addition, we hypothesize that longer genes have a larger number of regulatory elements that alter expression and enhance differential usage of these alternative sites. Long genes likely elongate during evolution, via insertions of TEs in their introns (Figure 7A,B; ***Sela et al., 2007; Grishkevich and Yanai, 2014).*** Long neuronal genes, such as ion channels and cell adhesion molecules, may be expressed primarily late in development ***(Okaty et al., 2009).*** Developmentally later and more spatially and cell-type restricted expression of neuronal genes may make mammalian genomes more tolerant to mutations caused by the insertion of TEs in these genes. Conversely, genes such as Hox genes, which are critical for early development, and are often expressed in progenitors giving rise to many cell types, are remarkably TE impoverished ***(Chinwalla et al., 2002; Simons, 2005).*** TE insertions occurring randomly are expected to happen more frequently in long genes (Figure 7F, Figure 7 Supplement 1F,G), thereby accelerating their elongation over time.

Here we provide evidence supporting the hypothesis that evolution of the vertebrate nervous system may have taken advantage of TE insertions and subsequent exaptations to diversify neuronal cell types, increasing the complexity of brain circuits. Long genes are enriched in the signaling molecules, receptors and ion channels responsible for input/output transformations in neurons, and the cell adhesion molecules that specify neuronal connectivity. Thus, changes in their expression could lead to changes in circuit level function. Specifically, elongation of long genes through TE insertions, occurring in the early embryo or in germ cells, likely creates a reservoir of genetic elements providing fodder for regulatory innovation. Subsequent exaptation of a fraction of these elements may have enhanced cell type-, and hence, behavioral-diversity, in turn, increasing the ability of populations to adapt to their environment (Figure 7F). This evolutionary advantage of lengthening neuronal genes may help to explain the paradox of why long genes should be abundantly expressed in CNS neurons despite the fact that these genes are sites of genome instability associated with genetic lesions leading to autism and other developmental disorders ***(Wei et al., 2016).*** This hypothesis also shifts focus away from short, developmental time scales considered in other hypotheses linking TE insertion to neuronal function ***(Muotri et al., 2005; Richardson et al.,2014; Perrat et al., 2013).*** Instead of DNA rearrangements in neuronal progenitors producing neuronal diversity, we consider the time scales of evolution and thus also shift focus to the germ line, where natural selection has its influence.

In summary, the elongation of neuronal effector genes may have endowed them with increased capacity for differential expression, permitting enhanced neuronal diversity. This diversity can also be characterized in terms of expression patterns of homeobox and other TFs. The maintenance of diverse neuronal identities must require interactions between expressed TFs and accessible ***cis*** regulatory elements within target effector genes. Identifying these interactions will require manipulating them within genetically identified cell types.

## Methods and Materials

### Cell Types and Mouse Lines

Cell types are defined operationally by the intersection of a transgenic mouse strain (or in some cases anatomical projection target) and a brain region. These “operational cell types” mayor may not correspond to “atomic” cell types, but as shown in Figure 2 have comparable purity to clusters of single cells. Mouse lines profiled in this study are summarized in Supplementary Table 1. Most were obtained from GENSAT ***(Gong et al., 2007)*** or from the Brandeis Enhancer Trap Collection ***(Shima et al., 2016).*** For Cre-driver lines, the Ai3, Ai9 or Ai14 reporter ***(Madisen et al., 2009)*** was crossed and offspring hemizygous for Cre and the reporter gene were used for profiling. All experiments were conducted in accordance with the requirements of the Institutional Animal Care and Use Committees at Janelia Research Campus and Brandeis University.

### Atlas

Animals were anesthetized and perfused with 4% paraformaldehyde and brains were sectioned at ***50μm*** thickness. Every fourth section was mounted on slides and imaged with a slide scanner equipped with a 20x objective lens (3DHISTECH; Budapest, Hungary). In house programs were used to adjust contrast and remove shading caused by uneven lighting. Images were converted to a zoomify compatible format for web delivery and are available at http://neuroseq.janelia.org.

### Cell Sorting

Manual cell sorting was performed as described ***(Hempel et al., 2007; Sugino et al., 2014).*** Briefly, animals were sacrificed following isoflurane anesthesia, and 300μm slices were digested with pronase E (1mg/ml, P5147; Sigma-Aldrich) for 1 hour at room temperature, in artificial cerebrospinal fluid (ACSF) containing 6,7-dinitroquinoxaline-2,3-dione ***(20μM***; Sigma-Aldrich), D-(-)-2-amino-5-phosphonovaleric acid (50μM; Sigma-Aldrich), and tetrodotoxin (0.1 μ***M***; Alomone Labs). Desired brain regions were micro-dissected and triturated with Pasteur pipettes of decreasing tip size. Dissociated cell suspensions were diluted 5-20 fold with filtered ACSF containing fetal bovine serum (1%; HyClone) and poured over Petri dishes coated with Sylgard (Dow Corning). For dim cells, Petri dishes with glass bottoms were used. Fluorescent cells were aspirated into a micropipette (tip diameter 30-50μm) under a fluorescent stereomicroscope (M165FC; Leica), and were washed 3 times by transferring to clean dishes. After the final wash, pure samples were aspirated in a small volume (1 *~3μl)* and lysed in *47μl* XB lysis buffer (Picopure Kit, KIT0204; ThermoFisher) in a *200μ1* PCR tube (Axygen), incubated for 30min at 40°C on a thermal cycler and then stored at -80°C. Detailed information on profiled samples are provided in Supplementary Table 2.

### RNA-seq

Total RNA was extracted using the Picopure kit (KIT0204; ThermoFisher). Either 1*μl* of 10^-5^ dilution of ERCC spike-in control (#4456740; Life Technologies) or (number of sorted cells/50) * (1 μ*l* of 10^-5^ dilution of ERCC) was added to the purified RNA and speed-vacuum concentrated down to *5μl* and immediately processed for reverse transcription using the NuGEN Ovation RNA-Seq System V2 (#7102; NuGEN) which yielded 4~8μg of amplified DNA. Amplified DNA was fragmented (Covaris E220) to an average of ~200bp and ligated to Illumina sequencing adaptors with the Encore Rapid Kit (0314; NuGEN). Libraries were quantified with a KAPA Library Quant Kit (KAPA Biosystems) and sequenced on an Illumina HiSeq 2500 with 4 to 32-fold multiplexing (single end, usually 100bp read length, see Supplemental Table 2).

### RNA-seq analysis

Adaptor sequences (AGATCGGAAGAGCACACGTCTGAACTCCAGTCAC for Illumina sequencing and CTTTGTGTTTGA for NuGEN SPIA) were removed from de-multiplexed FASTQ data using cutadapt v1.7.1 (http://dx.doi.org/10.14806/ej.17.1.200) with parameters “-overlap=7 -minimum-length=30". Abundant sequences (ribosomal RNA, mitochondrial, Illumina phiX and low complexity sequences) were detected using bowtie2 ***(Langmead and Salzberg, 2012)*** v2.1.0 with default parameters. The remaining reads were mapped to the UCSC mm10 genome using STAR ***(Dobin et al., 2012)*** v2.4.0i with parameters “-chimSegmentMin 15 -outFilterMismatchNmax 3”. Mapped reads are quantified with HTSeq ***(Anders et al., 2014)*** using Gencode.vM13 ***(Harrow et al., 2012).***

### Pan-neuronal genes

Pan-neuronal genes are extracted as satisfying the following conditions: 1) mean neuronal expression level (NE)> 20 FPKM, 2) minimum NE > 5 FPKM, 3) mean NE > maximum non-neuronal expression level (NNE), 4) minimum NE > mean NNE, 5) mean NE > 4x mean NNE, 6) mean NE > mean NNE + 2x standard deviation of NNE, 7) mean NE - 2x standard deviation of NE > mean NNE.

### DI/SC/DN calculation

To calculate DI, the following criteria were used to assign a “1” or “0” to each element in the difference matrix (DM): log fold change > 2 and q-value <0.05. Q-values were calculated using the limma package including the voom method (***Law et al., 2014).*** To adjust the power to be similar across cell types, two replicates (the most recent two) are used for all cell types with more than two replicates. We have tried the same calculations with 3 replicates (using a fewer number of cell types) and obtained similar results (data not shown).

To calculate binary DI (bDI), the following DM criteria were used: expression levels of all the replicates in one of the cell types in the pair < 1FPKM and expression levels of all the replicates in the other cell type in the pair > 15FPKM, in addition to q-value <0.05.

To assess the extent of differentiation by alternative splicing, we calculate differentiation at the level of each splice branch. See Figure 3D for the definitions of a splice branch and of branch probability. For each branch, at each alternative splice site, we define each pair of cell types as “different” when 1) branch probabilities for all replicates in a group are less than 0.3 or greater than 0.7, and 2) both cell types in the pair have > 10 reads reads at the alternative site. Condition 1) is justified by the bimodal distribution of branch probabilities shown in Figure 3E. Accumulating over all pairs creates a DM for each branch. We then combine all the branches using a logical “OR” to create a gene-level DM for each gene. If any branch distinguishes a pair of cell types, that pair is called “different” at the gene level. The gene-level DM has a value of “1” for pairs of cell types distinguished by any of the branches belonging to that gene, and has a value of “0” for pairs of cell types not distinguished by any branch belonging to the gene. The number of pairs compared can differ, depending on the expression pattern of the gene, since branch probabilities can only be calculated for cell types that express the gene. This situation differs from that for DI or bDI (based on expression levels rather than splicing) since pairs of cell types can be distinguished even if one does not express the gene. Therefore, unlike DI and bDI which assume a fixed number of total pairs, we use DN (total number of pairs distinguished), rather than the fraction of pairs distinguished, to rank genes.

### NNLS/Random forest decomposition

SCRS datasets deposited in NCBI GEO (GSE71585, **Tasic et al. (2016);** and GSE60361, **Zeisel et al. (2015))** were used for NNLS decomposition. Specifically the deposited count data were converted to TPM and used for comparison. The NeuroSeq dataset was quantified using RefSeq and featurecount ***(Liao et al., 2013)*** and converted into TPM. Subsets of genes common to all three datasets are then used for all further analyses. Since distributions of TPM values differed between datasets, they were quantile normalized to an average profile generated from the NeuroSeq dataset. Since most genes in the SCRS profiles exhibited noisy expression patterns, using the entire gene set for decomposition is not feasible. Therefore, we selected for decomposition the genes deemed most informative for distinguishing cell classes based on ANOVA across cell classes. However, simply taking the top ANOVA genes lead to highly biased gene selection since some cell types exhibited much larger transcriptional differences than others (e.g. many ANOVE selected genes were specific to microglia). We therefore selected genes so as to minimize the overlap between the cell types distinguished. Beginning with the highest ANOVA gene (highest ANOVA F-value), genes were selected only if their DM (Differentiation Matrix defined in Figure 3) differed from those previously selected, defined with a Jaccard index threshold of 0.5. We chose 300 genes from each dataset, yielding a total of 563 genes when all three sets were combined. This gene set was then used for all decompositions. Decompositions were performed on average profiles created by summing NeuroSeq replicates or by summing single-cell profiles using cluster assignments provided by the authors. NNLS was implemented using the Python scipy library (http://www.scipy.org).

For Random forest, implementation in the Python scikit-learn library ***(Pedregosa et al., 2011)*** was used.

### ATAC-seq

7 cell types, Purkinje and granule cells from cerebellum, excitatory layer 5, 6 and entorhinal pyramidal cells from cortex, excitatory CA1, or CA1-3 pyramidal cells from hippocampus, labeled in mouse lines P036, P033, P078, 56L, P038, P064, and P036 respectively (all from **Shima et al., 2016)** were profiled with ATAC-seq. They were FACS sorted to obtain ~20,000 labeled neurons. ATAC libraries for Illumina next-generation sequencing were prepared in accordance with a published protocol ***(Buenrostro et al.,2013).*** Briefly, collected cells were lysed in buffer containing 0.1% IGEPAL CA-630 (I8896, Sigma-Aldrich) and nuclei pelleted for resuspension in tagmentation DNA buffer with Tn5 (FC-121-1030, Illumina). Nuclei were incubated for 20-30 min at 37°C. Library amplification was monitored by real-time PCR and stopped prior to saturation (typically 8-10 cycles). Library quality was assessed prior to sequencing using BioAnalyzer estimates of fragment size distributions looking for a ladder pattern indicative of fragmentation at nucleosome intervals as well as qPCR to determine relative enrichment at two housekeeping genes compared to background (specifically the TSS of *Gapdh* and *Actb* were assessed relative to the average of three intergenic regions). For sequencing, Illumina HiSeq 2500 with 2 to 4-fold multiplexing and paired end 100bp read length was used. In addition to ATAC-seq, RNA-seq was performed on replicate samples of ~2,000 cells collected in a similar way, and library prepared using the same method described above.

### ATAC-seq analysis

Nextera adaptors (CTGTCTCTTATACACATCT) were trimmed from both ends from de-multiplexed FASTQ files using cutadapt with parameters “-n 3 -q 30,30 -m 36”. Reads were then mapped to UCSC mm10 genome using bowtie2 ***(Langmead and Salzberg, 2012)*** with parameters “-X2000 -no-mixed - no-discordant”. PCR duplicates were removed using Picard tools (http://broadinstitute.github.io/picard, v2.8.1) and reads mapping to mitochondrial DNA, scaffolds, and alternate loci were discarded. BigWig genomic coverage files were generated using bedtools ***(Quinlan and Hall, 2010)*** and scaled by the total number of reads per million. For reproducible peaks, liberal peaks were called using HOMER (v4.8.3) ***(Heinz et al., 2010)*** with parameters “-style factor -region -size 90 -fragLength 90 -minDist 50 -tbp 0 -L 2 -localSize 5000 -fdr 0.5” and filtered using the Irreproducibility Discovery Rate (IDR) in homer-idr (http://github.com/karmel/homer-idr.git) with parameters “-threshold 0.05 -pooled-threshold 0.0125”. Peak counts and peak patterns were then quantified using bedtools.

### TF Tree

The set of mouse TFs was constructed by combining 4 curated TF lists: genes annotated in 1) PANTHER ***(Thomas, 2003)*** PC00218 (transcription factor), 2) Riken Transcription Factor Database ***(Kanamori et al., 2004),*** 3) HUGO ***(Grayetal., 2014)*** families with TF functions and 4) Gene Ontology ***(Ashburner et al., 2000***) G0:0006355 (regulation of transcription). Genes appearing reproducibly in these list (i.e. in more than 1 list) were used as TFs. Anatomical regions used as constraints are defined in a hierarchical manner (see Supplementary Table 5).

The TF tree is constructed recursively using the following algorithm:

#### preparation

1. `calculate bDIs for all subsets of samples defined by anatomical regions function bisect(list of samples):`
2. `if the list of samples consists of only one cell type, exit`
3. `calculate bDI,SC within this group of samples for all TFs`
4. `if there is no TF with bDI>0, exit`
5. `find the appropriate level in the hierarchy of anatomical regions`
6. `penalize bDIs (from 2.) with bDIs of containing anatomical regions (from 0.)`
7. `sort TFs by their penalized bDI and SC in descending order`
8. `set candidates as TFs with penalized bDI>0.2, if there are none, take the top 5`
9. `for each candidate, calculate divisions of samples according to expression level`
  `- at sample level, assign ON/OFF using FPKM=3 as threshold`
  `- at cell type level, assign ON/OFF according to dominant ON/OFF of samples`
  `- divide all cell types into ON or OFF groups`
  `- optionally constrain division to anatomical boundary`
10. `if there is no division, exit`
11. `if there is more than one division then`
  `-calculate “division strength” for all divisions:`
    `-a0 = mean number of binary distinctions of all genes between ON and OFF groups`
    `-a1 = mean number of binary distinctions of all genes within ON or OFF groups`
    `-division strength = a0/a1`
  `-then choose the division with the highest division strength`
12. `output ON/OFF groups and corresponding TF(s) for the chosen division`
13. `call bisect on ON group samples`
14. `call bisect on OFF group samples`

### Inserted segments

The multiz alignments downloaded from the UCSC genome browser ***(Kent et al., 2002)*** was used to calculate inserted segments in human or mouse. By comparing closely related species (human vs. chimp or mouse vs. rat), candidate segments inserted into human (or mouse) are extracted.By using another closely related species as a common ancestor (gorilla, guinea pig respectively for human/chimp and mouse/rat), segments absent in chimp and gorilla (or absent in rat/guinea pig) are called insertion in human (or mouse), and segments absent in chimp but present in gorilla (or absent in rat but present in guinea pig) are called deletion in chimp (or rat).

### TE fitting

Repeat annotations for mouse mm10 genome as detected by RepeatMasker ***(Smit et al., 2013-2015)*** with Repbase (ver. 20140131 **Bao et al., 2015)** were used. Only repeat families with number of instances>200 are included. For individual repeats, only those with number of instances>50 are included. For repeats in the “Simple repeat” class, only those with number of instances>1000 are included. Repeat scores are calculated as described in Figure 7D using Gencode.vM13. Only genes with non-zero repeat scores are used for fitting. For fitting expression level (rank) by repeat score, a regularized version of linear regression, Ridge regression, was implemented in the Python scikit-learn library ***(Pedregosa et al., 2011).***

### Tissue data

In addition to cell type-specific data obtained in this study, we analyzed publicly available RNA-seq and DNase-seq data using tissue samples. Information on these samples are described in Supplementary Table 4.

### Annotations

For reference annotations we used Gencode.vM13 ***(Harrow et al., 2012)*** downloaded from http://www.gencod NCBI RefSeq ***(Pruitt et al., 2013)*** downloaded from the UCSC genome browser.

### Anatomical Region Abbreviations

Region abbreviations: AOBmi, Accessory olfactory bulb, mitral layer; MOBgl, Main olfactory bulb, glomerular layer; PIR, Piriform area; COAp, Cortical amygdalar area, posterior part; AOBgr, Accessory olfactory bulb, granular layer; MOBgr, Main olfactory bulb, granular layer; MOBmi, Main olfactory bulb, mitral layer; VISp, Primary visual area; AI, Agranular insular area; MOp5, Primary motor area, layer5; VISp6a, Primary visual area, layer 6a; SSp, Primary somatosensory area; SSs, Supplemental somatosensory area; ECT, Ectorhinal area; ORBm, Orbital area, medial part; RSPv, Retrosplenial area, ventral part; ACB, Nucleus accumbens; OT, Olfactory tubercle; CEAm, Central amygdalar nucleus, medial part; CEAl, Central amygdalar nucleus, lateral part; islm, Major island of Calleja; isl, Islands of Calleja; CP, Caudoputamen; CA3, Hippocampus field CA3; DG, Hippocampus dentate gyrus; CA1, Hippocampus field CA1; CA1sp, Hippocampus field CA1, pyramidal layer; SUBd-sp, Subiculum, dorsal part, pyramidal layer; IG, Induseum griseum; CA, Hippocampus Ammon's horn; PVT, Paraventricular nucleus of the thalamus; CL, Central lateral nucleus of the thalamus; AMd, Anteromedial nucleus, dorsal part; LGd, Dorsal part of the lateral geniculate complex; PCN, Paracentral nucleus; AV, Anteroventral nucleus of thalamus; VPM, Ventral posteromedial nucleus of the thalamus; AD, Anterodorsal nucleus; RT, Reticular nucleus of the thalamus; MM, Medial mammillary nucleus; PVH, Paraventricular hypothalamic nucleus; PVHp, Paraventricular hypothalamic nucleus, parvicellular division; SO, Supraoptic nucleus; DMHp, Dorsomedial nucleus of the hypothalamus, posterior part; ARH, Arcuate hypothalamic nucleus; PVHd, Paraventricular hypothalamic nucleus, descending division; SCH, Suprachiasmatic nucleus; LHA, Lateral hypothalamic area; SFO, Subfornical organ;

VTA, Ventral tegmental area; SNc, Substantia nigra, compact part; SCm, Superior colliculus, motor related; IC, Ingerior colliculus; DR, Dorsal nucleus raphe; PAG, Periaqueductal gray; PBl, Parabrachial nucleus, lateral division; PG, Pontine gray; LC, Locus ceruleus; CSm, Superior central nucleus raphe, medial part; AP, Area postrema; NTS, Nucleus of the solitary tract; MV, Medial vestibular nucleus;NTSge, Nucleus of the solitary tract, gelatinous part; DCO, Dorsal cochlear nucleus; NTSm, Nucleus of the solitary tract, medial part; IO, Inferior olivary complex; VII, Facial motor nucleus; DMX, Dorsal motor nucleus of the vagus nerve; RPA, Nucleus raphe pallidus; PRP, Nucleus prepositus; CUL4,5mo, Cerebellum lobules IV-V, molecular layer; CUL4,5pu, Cerebellum lobules IV-V, Purkinje layer; PYRpu, Cerebellum Pyramus (VIII), Purkinje layer; CUL4,5gr, Cerebellum lobules IV-V, granular layer; MOE, main olfactory epithelium; VNO, vemoronasal organ.

## Acknowledgments

We thank Jody Clements and Charlotte Weaver for help in preparing web site, Erina Hara, Asish Gulati, Xiaotang Jing and Zhe Meng for technical help, Keven McGowan for assistance in sequencing,Jim Cox, Amanda Zeladonis and Amanda Wardlaw for help in animal maintenance, Gabe Murphy for help in retinal sample collection, and Rosa Miyares for comments for the manuscript.

---

**Figure 1-Supplement 1.**
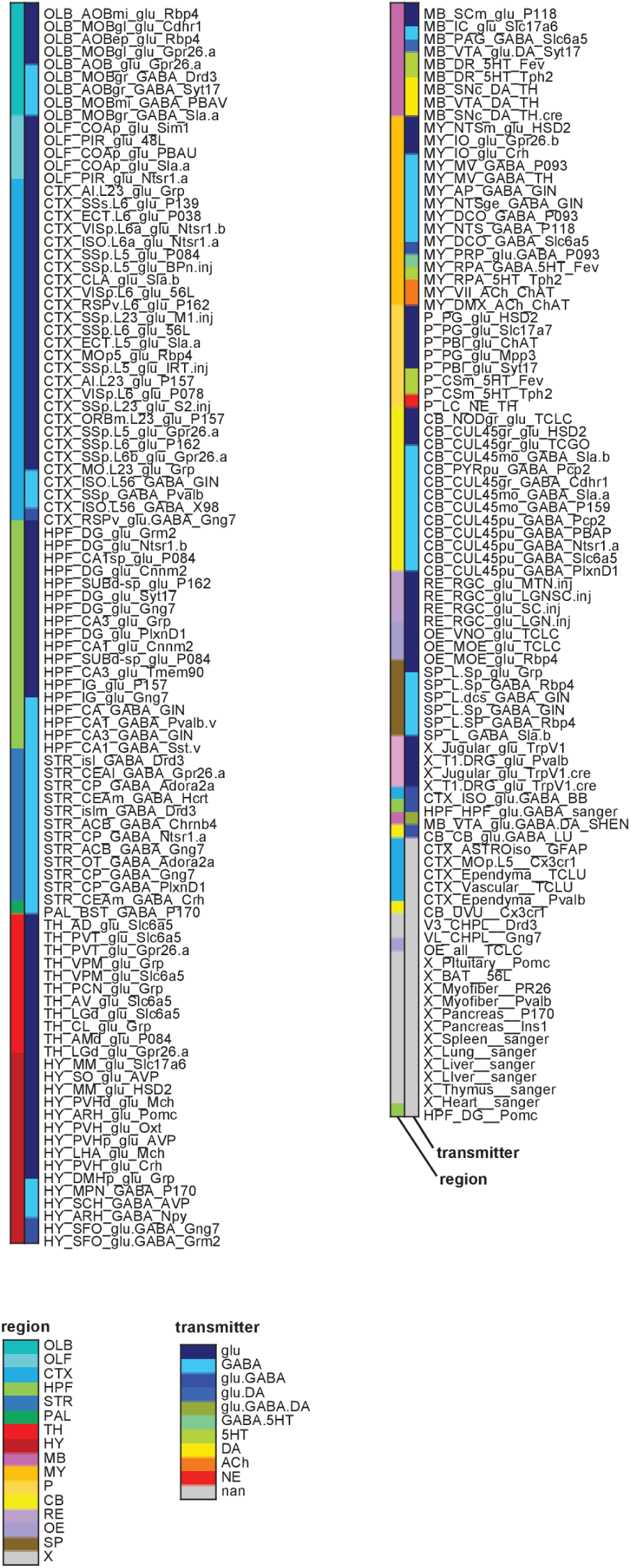
Cell type-specific samples. Sample groups color coded by region (left color bar) and transmitter phenotype (right color bar). Transmitter phenotype was determined from transmitter synthesis and storage enzyme expression. Abbreviations: OLB: olfactory bulb;OLF: olfactory regions (excluding bulb);CTX: Isocortex and Claustrum;HPF: hippocampal formation;STR: Striatum and related ventral forebrain structures;PAL: pallidum;TH: thalamus;HY: hypothalamus;MB: midbrain;MY: medulla; P: pons;CB: cerebellum;RE: retina;OE: olfactory epithelium;SP: spinal cord;X: peripheral nervous system or non-neural tissue. For additional abbreviations see Methods.

**Figure 1-Supplement 2.**
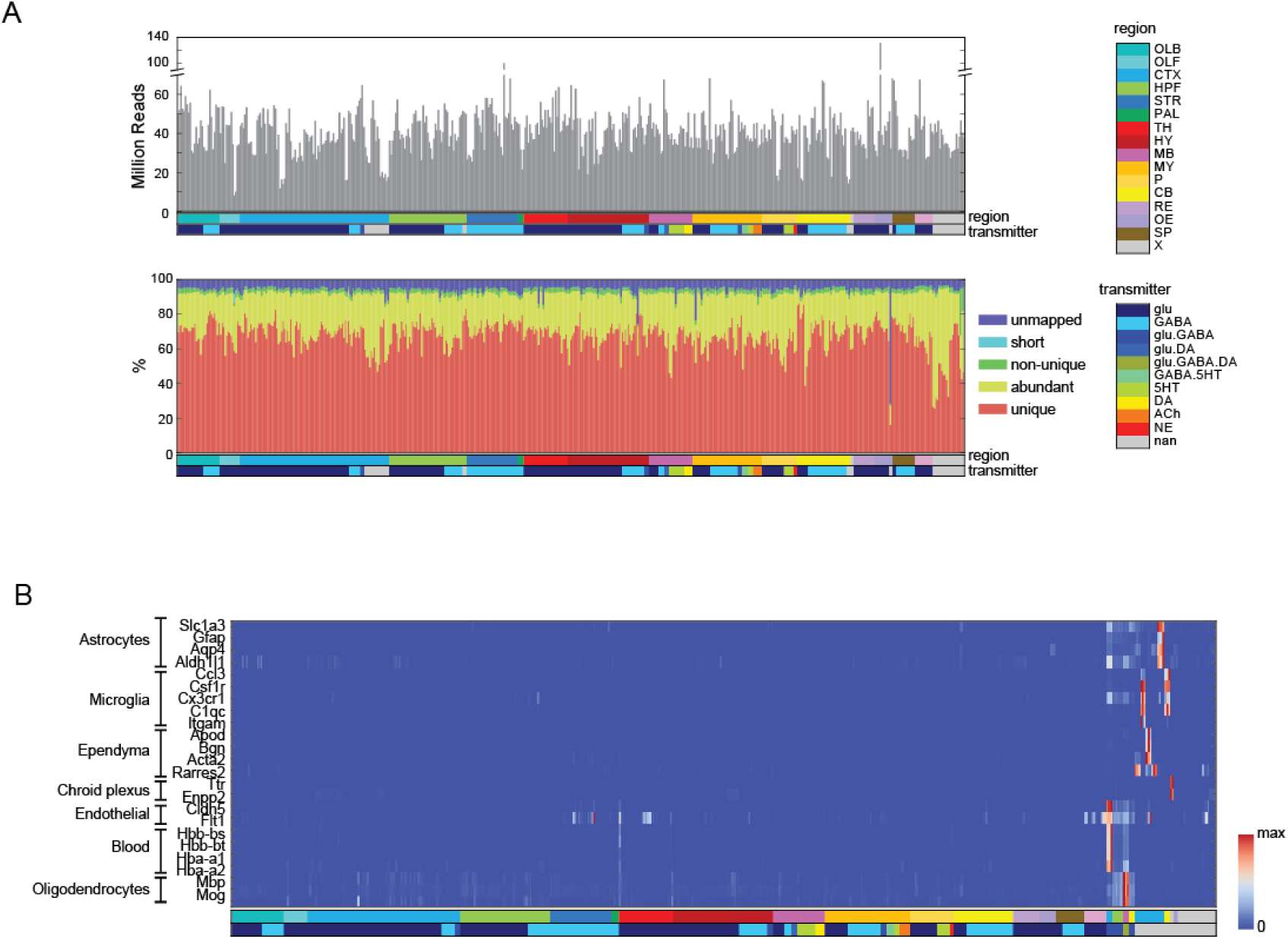
Quality Control measures. **(A)**(Top) Total reads for each of the libraries. Samples are color coded by region and transmitter, as shown in Figure 1 Supplement 1. (Bottom) Categories of reads in each library: unmapped: reads that did not map to the mm10 genome including chimeric and back-spliced reads; short: reads less than 30bp in length after removing adaptor sequences; non-unique: reads mapping to multiple locations; abundant: reads containing ribosomal RNA polyA, polyC and phiX sequences, and unique: uniquely mapped reads. For further analyses, abundant, short and unmapped reads were not used. **(B)** Contaminating transcripts from non-neuronal cell types. Samples with significant expression of these transcripts (at right) include tissue samples and non-neuronal samples. Each row is normalized by the maximum value.

**Figure 1-Supplement 3.**
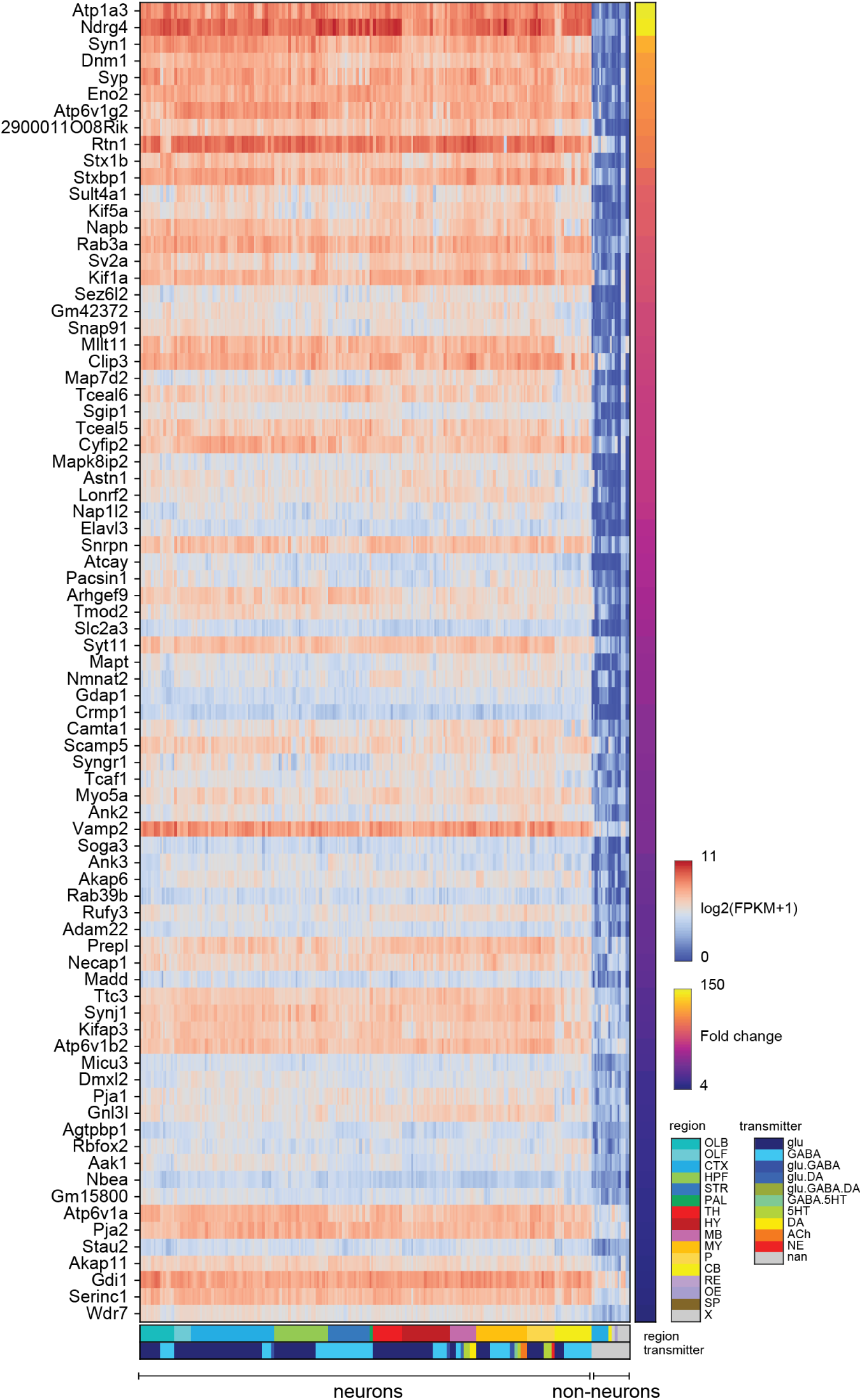
Pan-neuronal genes. Genes expressed in all neuronal cell types, but not (or at much lower levels) in non-neurons within the dataset. Heat-map shows log expression levels and the color at the right side indicates fold-change of the expression level between neurons and non-neurons. Criteria for extracting these genes are listed in the Methods.

**Figure 2-Supplement 1.**
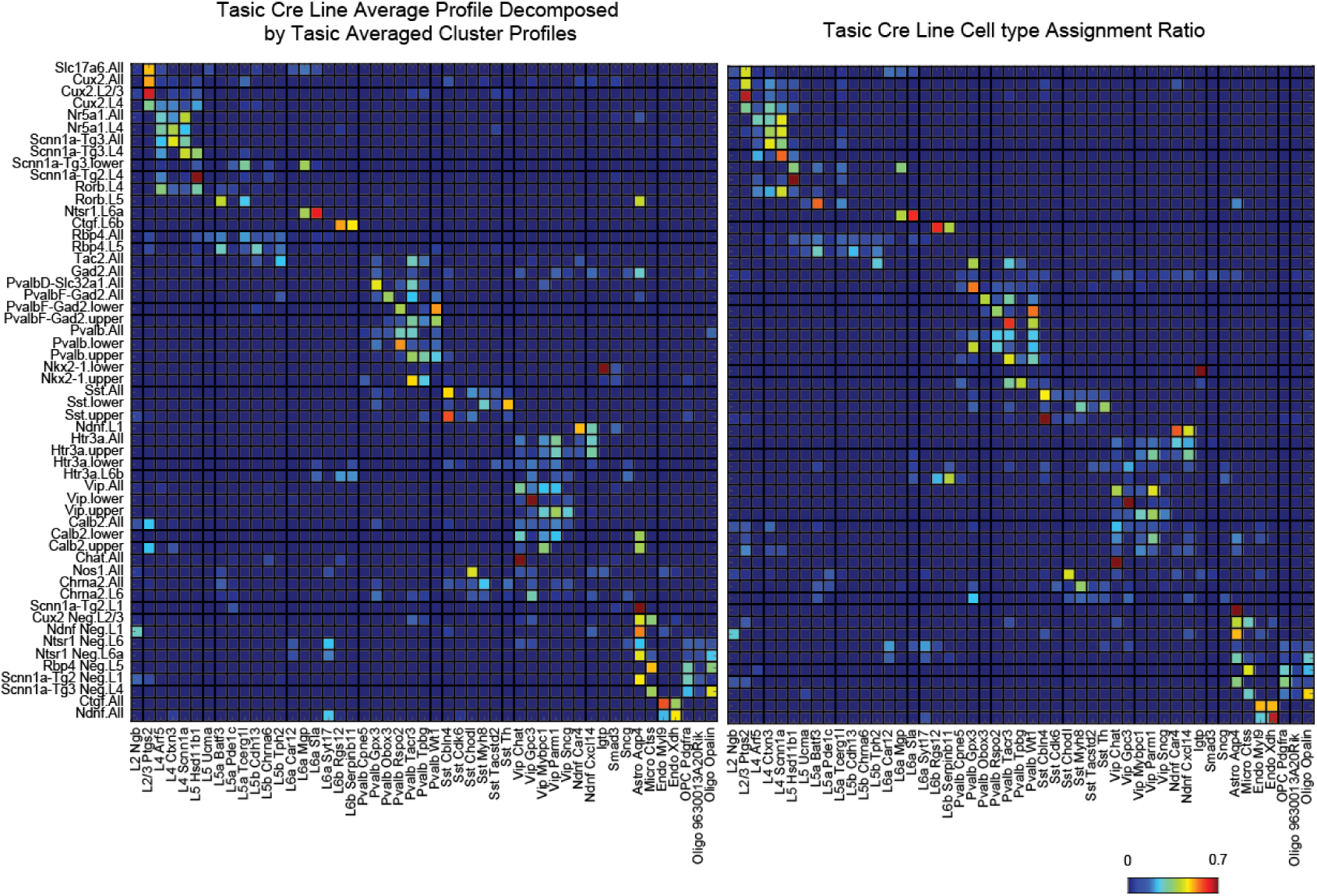
A test of NNLS decomposition. (Left) Single cell profiles from **Tasic et al. (2016)** were merged according to which of the 17 transgenic strains and sub-dissected layers they originated from (row labels). Merged profiles were then decomposed using NNLS by the same individual cluster profiles used in Figure 2 (column labels). **(Right)** The reported proportion of single cell profiles according to the author's classification. The close similarity between left and right matrices indicates an accurate NNLS decomposition of the merged clusters. Note that information about which and how many individual cell types were sorted from each line and set of layers was not explicitly provided to the decomposition algorithm, but were accurately deduced from the merged expression profiles.

**Figure 2-Supplement 2.**
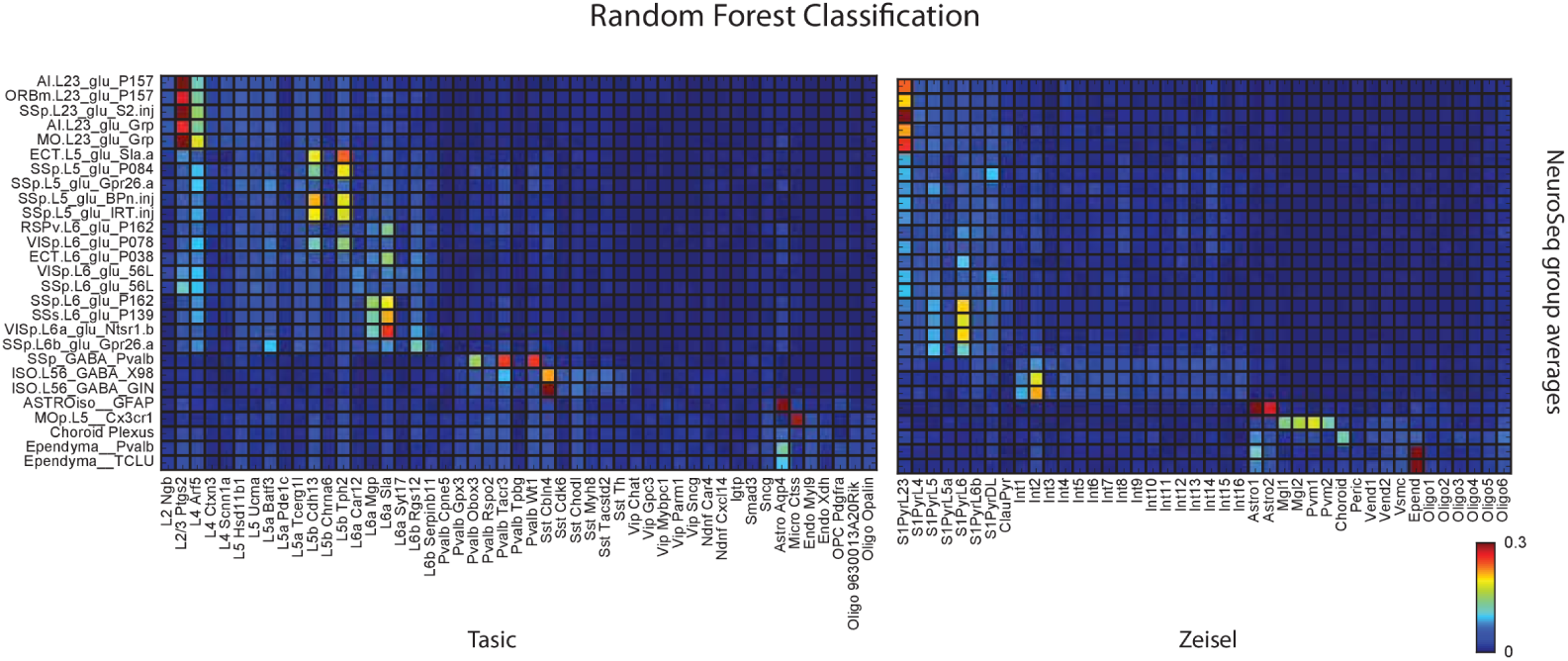
Random forest decomposition. A random forest classifier (500 decision trees) was trained from single cell profiles and their cluster assignment (column labels) and then used to decompose NeuroSeq cell types (row labels). Coefficients are the ratio of the votes from the 500 trees (coefficient ranges from 0 to 1 and 1 indicates all trees vote for a single class). The pattern of coefficients is similar to that obtained by NNLS (Figure 2A) suggesting the decomposition is relatively robust and does not reflect a peculiarity of the NNLS algorithm.

**Figure 2-Supplement 3.**
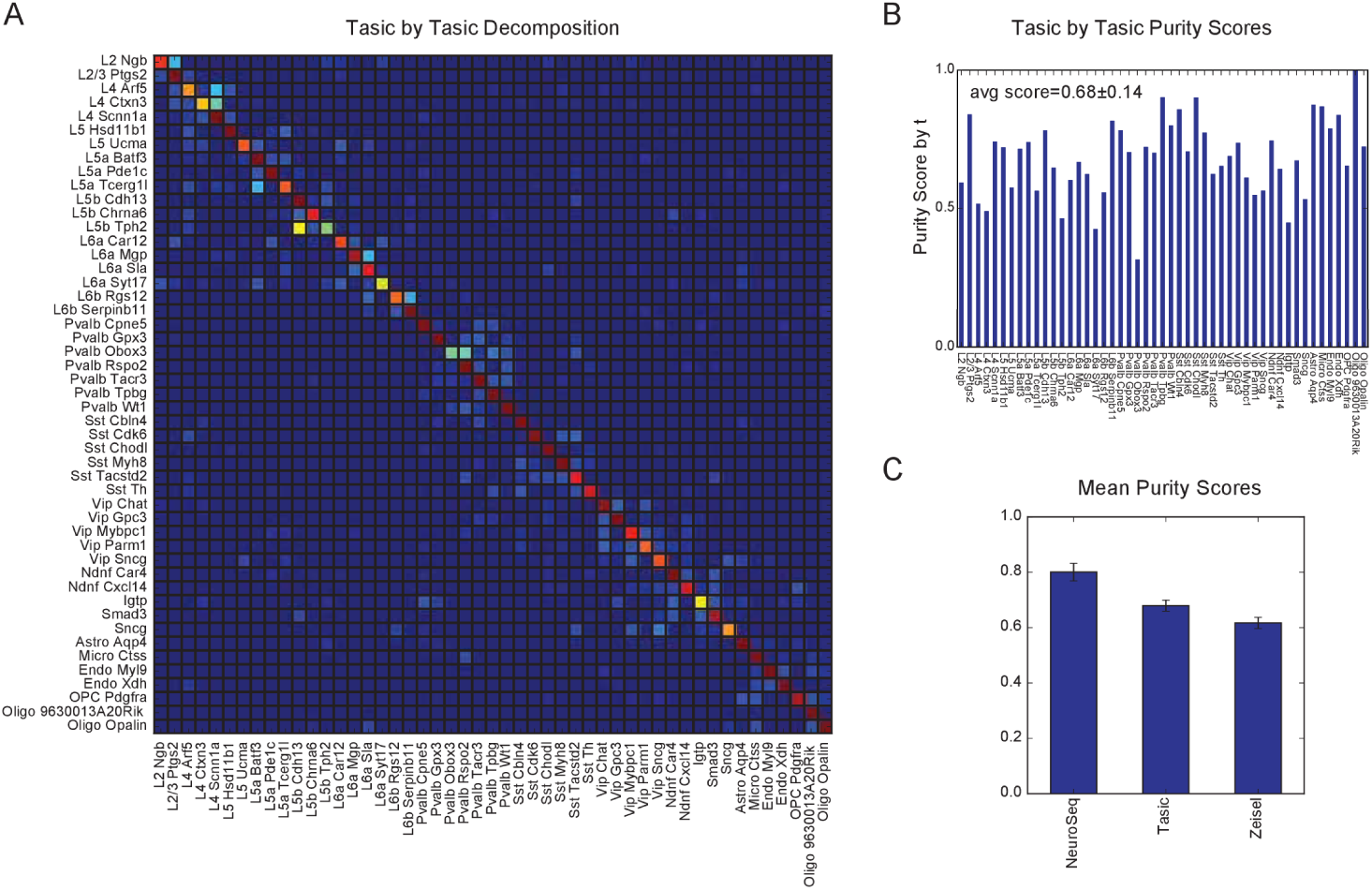
Cross validation of NNLS decompositions. **(A)** Each of Tasic et al. cluster is randomly divided into two groups and one is used to decompose the other. Some cluster pairs share significant coefficients, suggesting they are too similar to each other to separate well. For example, pairs of clusters L2 Ngb and L2/3 Ptgs2, L4 Arf5 and L4 Scnn1a, L4 Ctxn3 and L4 Scnn1a, and L5 Cdh13 and L5 Tph2 are hard to distinguish. This is consistent with the observation of intermediate cells between each of these clusters in the original study (their Figure 4).**(B)** Purity scores (similar to Figure 2C) for the cross-validated NNLS decomposition of each Tasic et al. cluster. **(C)** Mean purity scores obtained from the same cross-validation procedure applied to each of the three datasets.

**Figure 2-Supplement 4.**
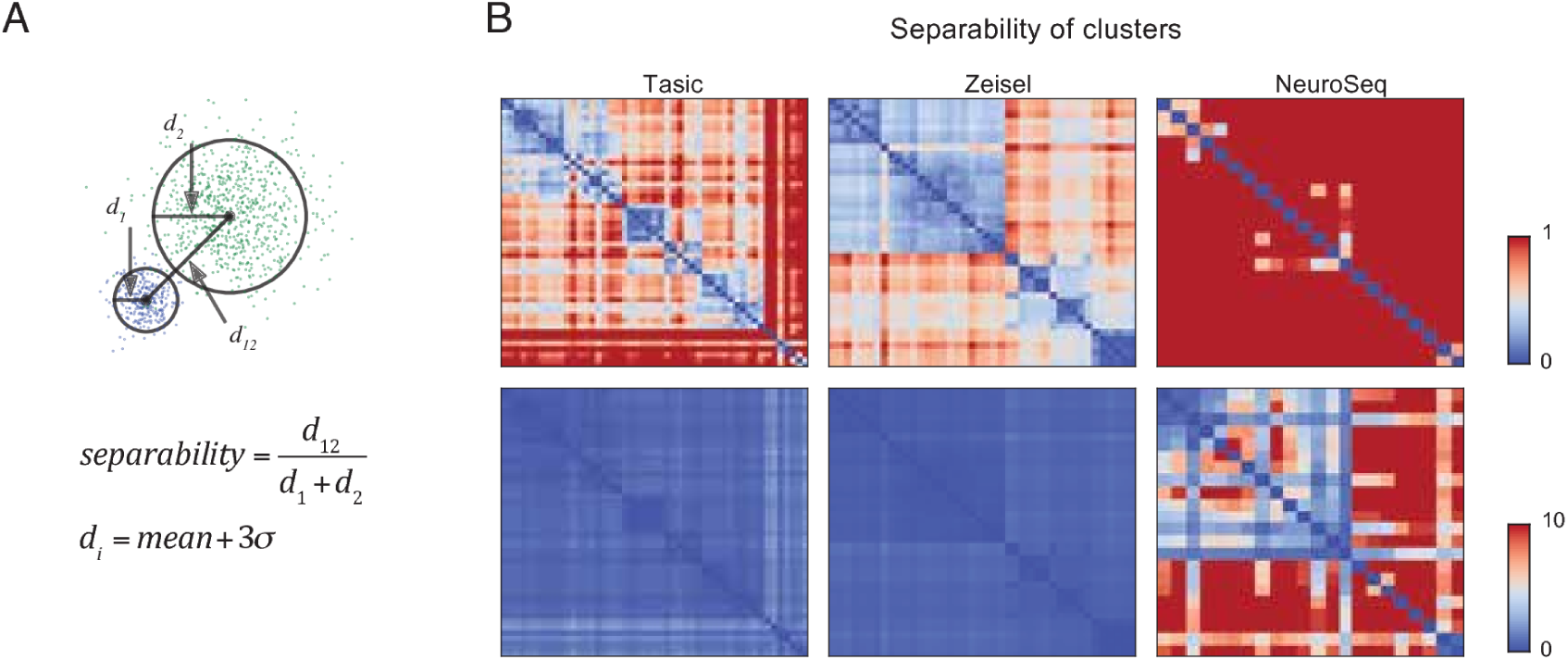
Separability of cell type clusters. **(A)** Definition of separability. Cartoon represents two different single cell clusters as distributions of points. The separability is the ratio of the distance between the centroids to the sum of the “diameter” of each cluster. Here, we calculate the diameter of a cluster using the distances from the centroid of the cluster as the mean distance + 3 times the standard deviation of the distribution of the distances. With this definition, two clusters are “touching” when separability =1, overlapping when <1, and separate when >1. The multi-dimensional distance is computed as 1-Pearson's corr.coef. **(B)** Separabilities between cell type clusters for three datasets shown with two different dynamic ranges (color scale;0-1 for upper row and 0-10 for lower row). The order of cell type clusters are the same as in Figure 2.

**Figure 3-Supplement 1.**
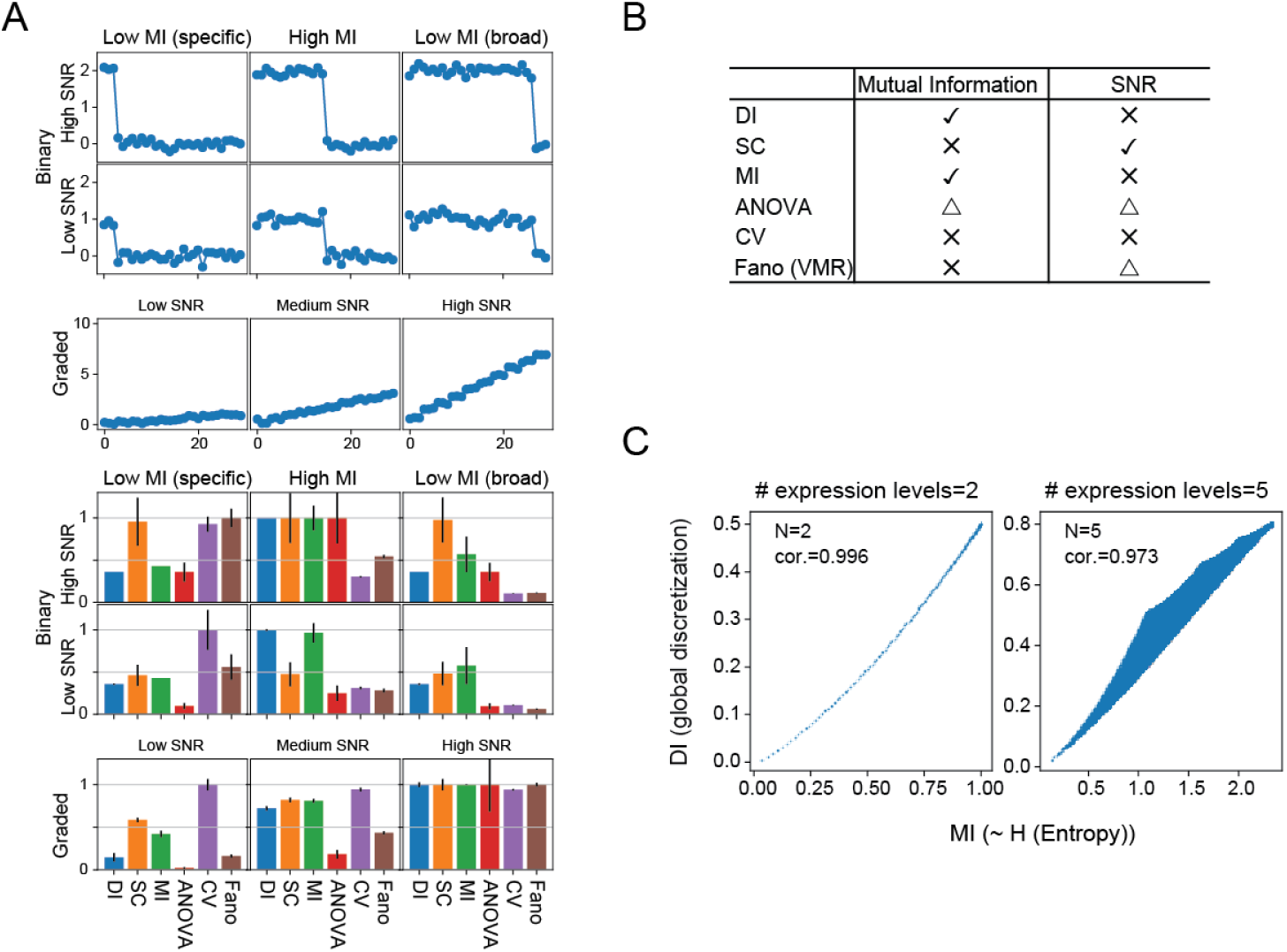
Simulated data reveal features of expression metrics. **(A)** (Upper) An example of simulated binary and graded expression patterns with added noise. X-axis indicates sample/groups. (Lower) Various average metrics calculated from the simulated expression patterns (100 individual simulations; error bars are standard deviations). Values are normalized within each metric across binary expression group or graded expression group. **(B)** Summary of each metric's correlation with Mutual Information and SNR: check mark-correlated, X-uncorrelated, triangle-partially correlated. **(C)** DI and MI are highly correlated. The relationship between DI, calculated without considering replicates, and MI with expression levels discretized into 2 levels (left) and 5 levels (right). Although increasing the number of discrete expression levels decreases the degree of correlation, they remain monotonically and closely related.

**Figure 3-Supplement 2. Relationship between DI and MI** Here we explore more detailed relationship between mutual information and differentiation index. To calculate mutual information between expression levels and cell types, we discretize expression levels into *N*_*e*_ levels. Let *N*_*s*_ be number of samples. Let *n*_*iJ*_ be counts in the contingency table where *i* = 1, *…,N*_*e*_ and *j =* 1, …,*N*_*S*_. Then the joint probability distribution and the marginal probability distribution can be written as:

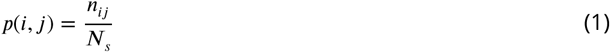

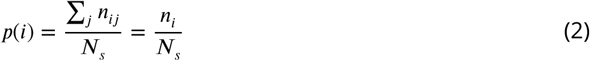

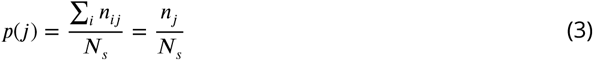

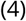

Where ***n_i_ =*** Σ_*j*_ *n*_***ij***_ and *n_j_ = Σ n*_*ij*_. *n*_*j*_ is number of replicates in cell type *j*. The mutual information between expression level (E) and samples (S) is:

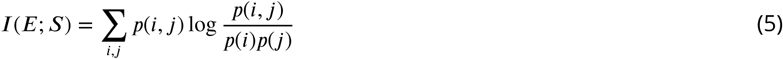

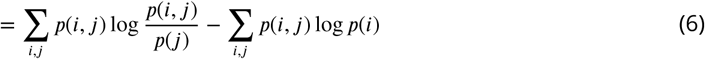

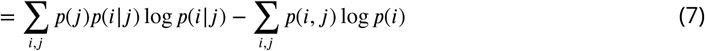

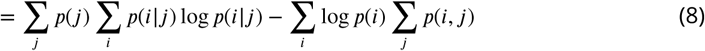

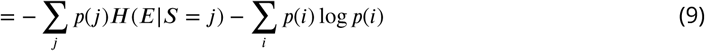

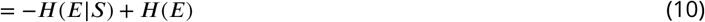

***H(E***∣***S*** = *j)* is the entropy of expression levels in cell type j, which represents the expression noise in cell type j, and ***H***(*E*∣*S*) is the average of these across all cell types. When there is no replicates ***H***(*E*∣*S*) is zero. When there are replicates, ***H***(*E*∣*S* = *j)* represents how noisy the expression is. This may depends on expression level, and ***H***(*E*∣*S*), the average of ***H***(*E*∣*S* = *j)* may depends on expression prevalence (i.e., how widely the gene is expressed), but in any case, the first term —***H(**E*∣*S*) represents reduction of the mutual information by noise.

The second term ***H(E)*** is the entropy of marginal distribution *p(i)* and represents the main information content of cell types encoded in expression levels. This can be rewritten using counts in the contingency table as:

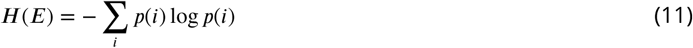

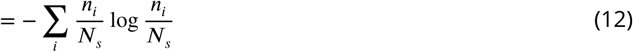

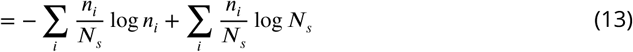

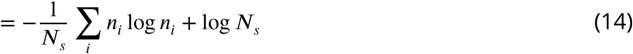

Thus, it takes maximum when all *n_i_’*s are 0 or 1, which corresponds to the case where one expression level corresponds to one cell type, making all cell types distinguishable by the expression levels. This is when the discretization levels are larger than number of samples. When the number of discretization levels (*N*_*e*_) is smaller than the number of samples (*N*_*s*_), *H(E)* takes the maximum value of log *N*_*e*_ when all the samples are distributed equally to each bin.

To explore the relationship between *H(E)* and DI, the log *n*_*i*_ in the first term is replaced (approximated) by ***(n_i_*** — 1) (first two terms in the Taylor expansion of log *n*_*i*_ around *n*_***i***_ = 1.):

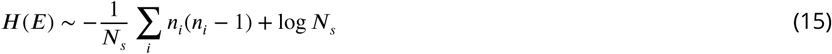

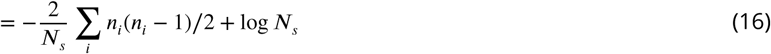

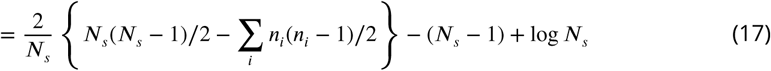

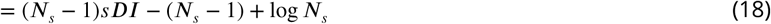

Since *n*_*i*_ is the number of samples in one expression level, *n*_*i*_(*n*_*i*_ — 1)/2 is the number of indistinguishable pairs in that expression level when there is no replicate. The term within the curly bracket is then the number of distinguishable pairs, leading to eq.(18).

More formally, since both *h(p) = Σn*_*i*_ log *n*_*i*_ and 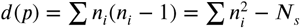 are Schur-convex functions^1^ on partitions of *N*_*s*_, *p =* (*n*_1_,*n*_2_,…,*n*_*k*_), when partition *p*_*1*_ majorizes *p*_*2*_ then, *h*(p_1_) ≥ *h(p*_2_) and *d(p*_1_) ≤ *d(p*_2_). When partition length is 2, that is when expression levels are discretized into only 2 levels, corresponding to ON/OFF, then, all of the partitions can be ordered by majorization relationship, therefore, *h(p)* and *d(p)* are order-preserved transformation of each other (Figure 3 Supplement 1C left). When partition length is greater than 2, this relationship is not true. However, they are still highly correlated to each other (Figure 3 Supplement 1C right).

When DI is calculated from global discretization (as in the above case), the maximum number of pairs distinguishable happens when all the samples are equally distributed to each bin and the number of distinguishable pairs is 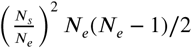. Therefore,

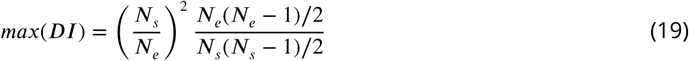

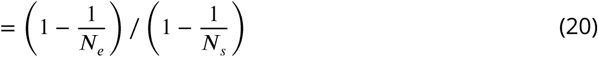

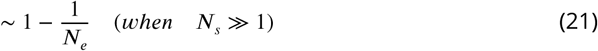

As stated above, this is also when the entropy ***H(E)*** takes the maximum value of log_2_ *N*_*e*_ in the unit of bits. (Figure 3 Supplement 1C)

**Figure 4-Supplement 1.**
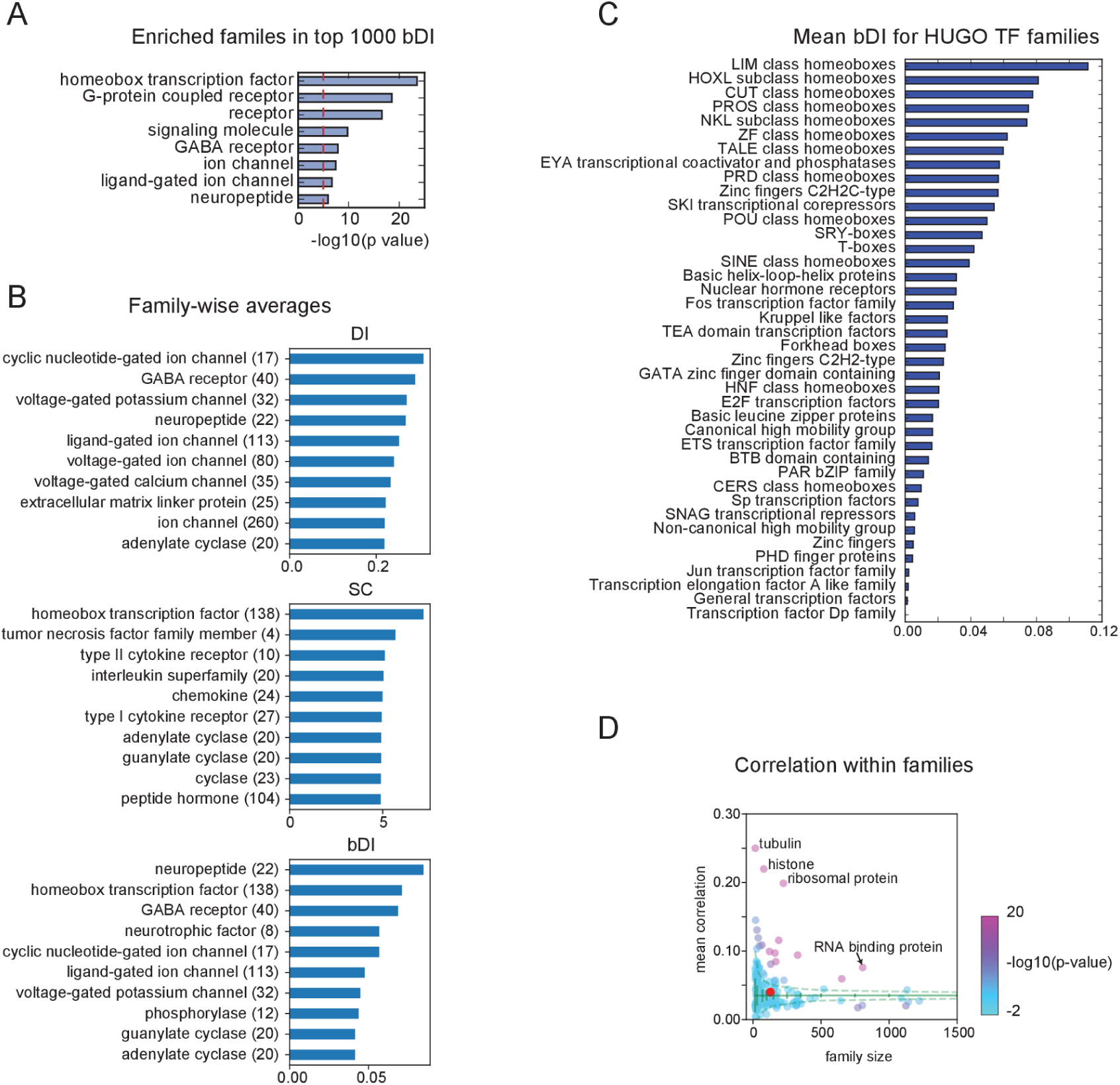
**(A)** PANTHER families enriched in the top 1000 bDI genes. **(B)** Averages of metrics (DI,SC,bDI) for PANTHER families. Only top 10 are shown. Numbers in parenthesis indicate family size. **(C)** Average bDI calculated for each TF family in HUGO protein families **(Gray et al., 2014). (D)** Mean Pearson's corr. coef. between genes within PANTHER families. Homeobox TF family is indicated by the red dot. Most of the PANTHER family genes are decorrelated within families but genes in some families, such as ribosomal protein, histone, tubulin, and RNA binding protein have highly significant correlation within families.

**Figure 5-Supplement 1.**
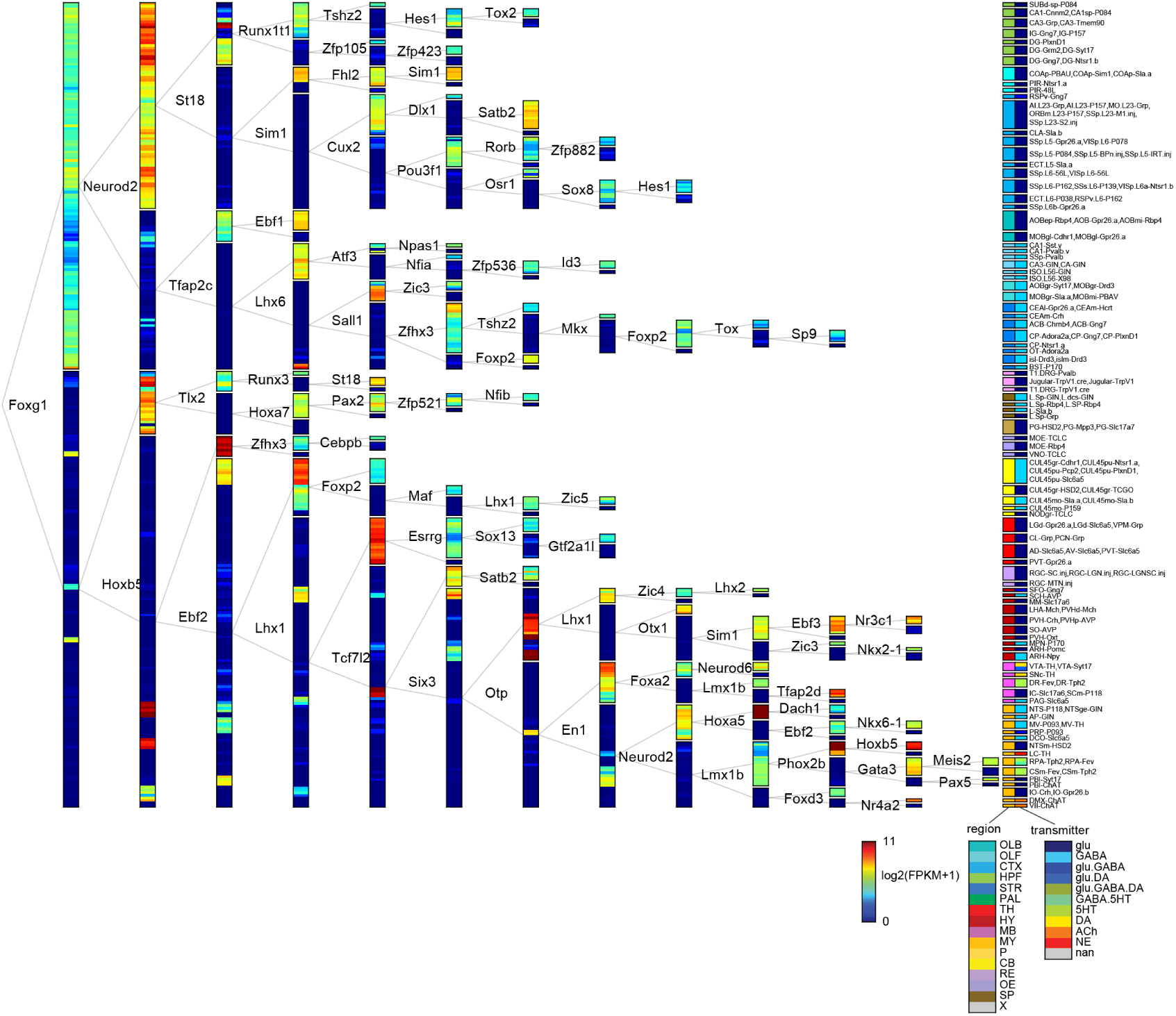
TF tree constructed using stronger anatomical constraints. Similar to Figure 5, but the constraints on anatomical boundaries are enforced during each bisection. However, TF expression was not constrained to be uniform within a group, leading to some subgroups that do not match the expression of the dividing gene.

**Figure 5-Supplement 2.**
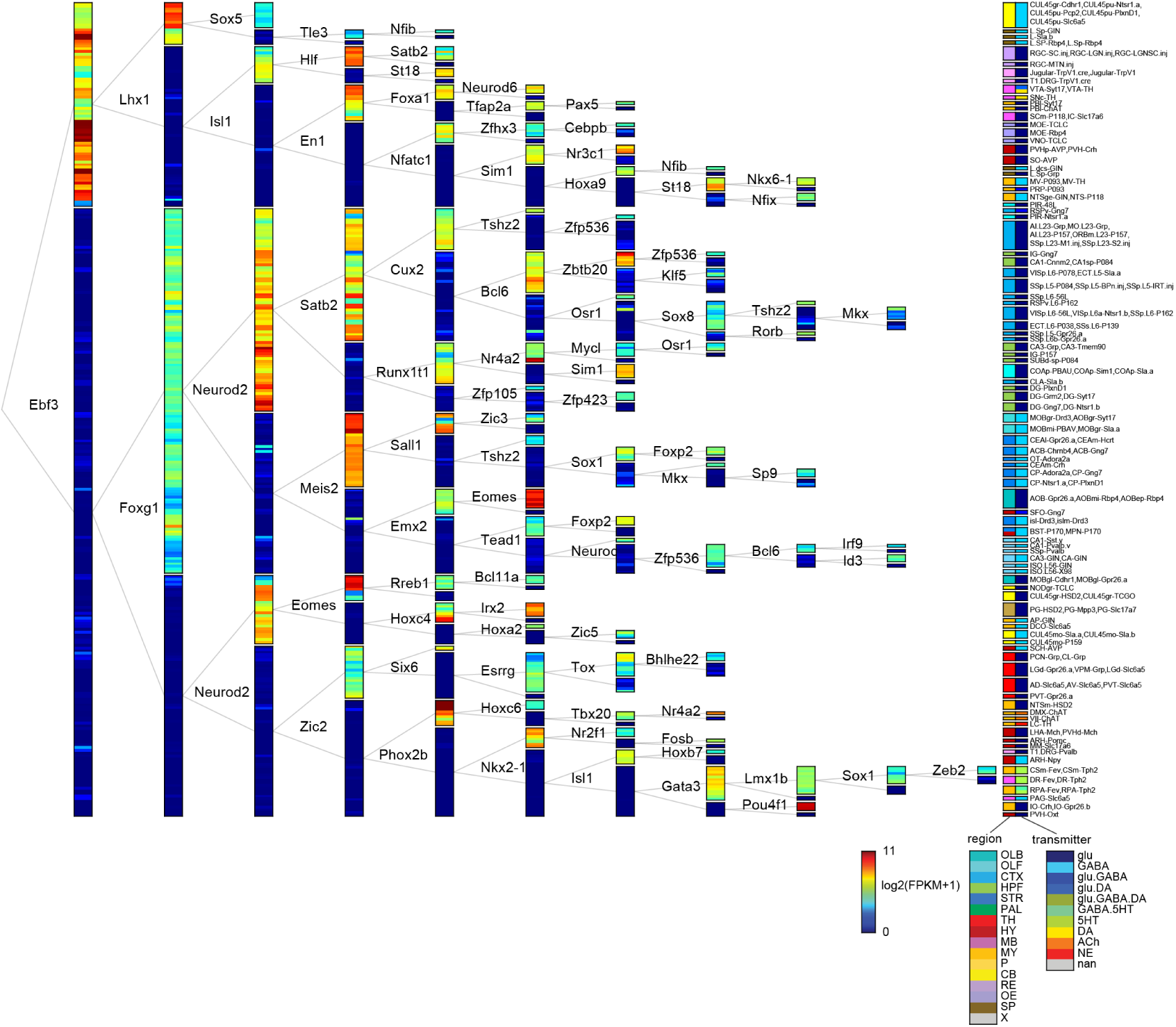
TF tree constructed without anatomical constraints. Similar to Figure 5 but anatomical subregions were not constrained to be grouped together.

**Figure 6-Supplement 1.**
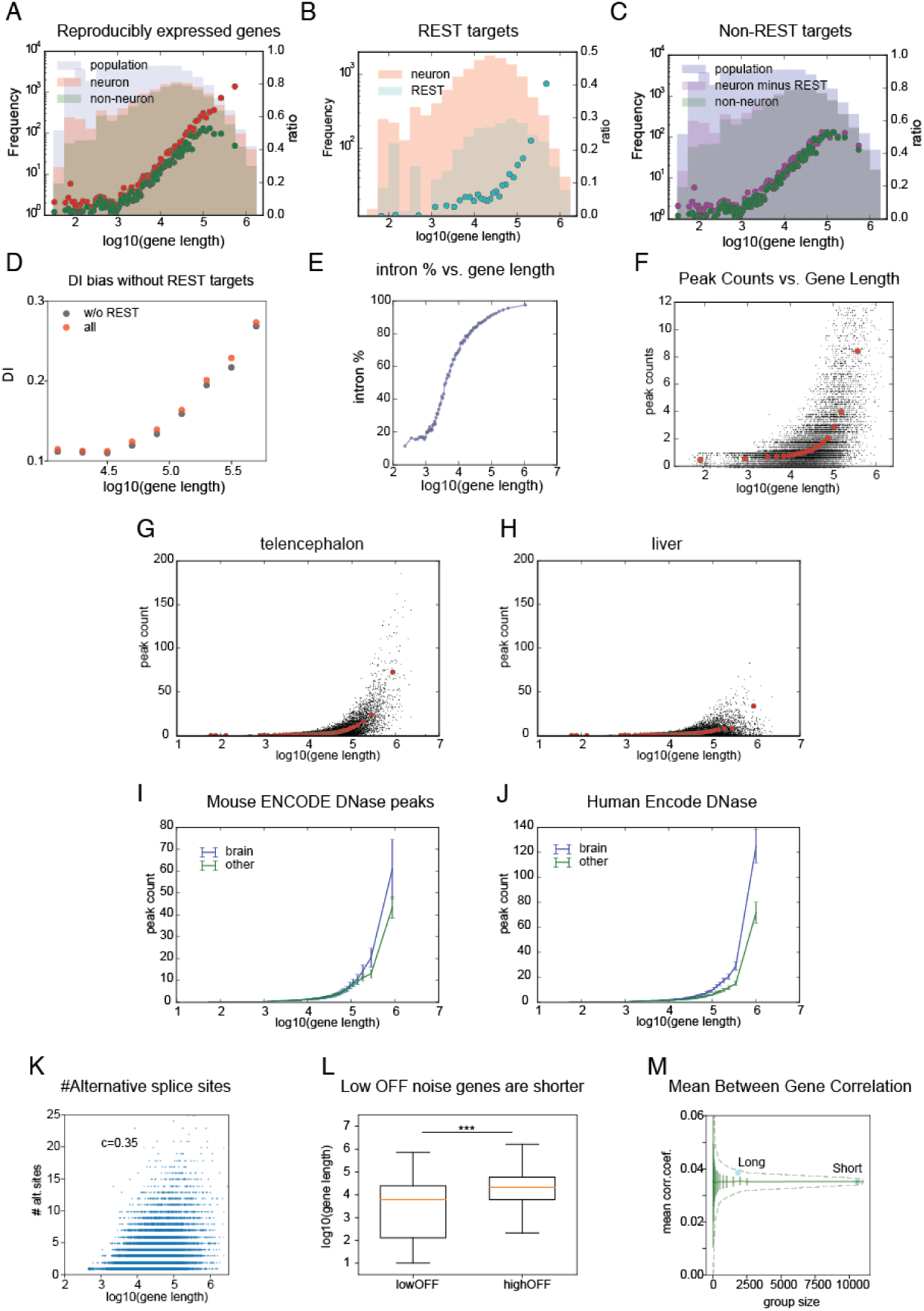
Properties of long genes in current and prior datasets. **(A)** Number (histogram) and ratios (dots) of genes expressed in neurons (pink histogram, red dots) and non-neurons (brown histogram, green dots) relative to the number of genes in the entire population (grey histogram) as a function of gene length (ratios computed per bin of 500 genes). ***(B)*** Number (cyan histogram; left axis) and ratios (cyan dots; right axis) of genes with nearby NRSE relative to the numbers of neuronally expressed genes (pink histogram). ***(C)***(Magenta dots) ratio of neuronally expressed non-REST target genes to the population. Other components are same as in A. ***(D)*** DI dependence of length without REST target genes compared to all genes. DI is still strongly length dependent because REST targets are a small fraction of expressed long genes. ***(E)*** Fraction of gene length attributable to intron length. ***(F)*** Length dependence of peak counts in the ATAC-seq data from the current study. ***(G)-(J)*** Length dependence of peak counts in ENCODE DNase hypersensitivity data. Examples from mouse ENCODE data in forebrain (telencephalon) ***(G)*** and liver ***(H)*** samples showing individual peaks (black dots) and binned averages (red dots) as a function of gene length. Average mouse ***(I)*** and human ***(J)*** peak counts from brain(blue) and non-brain(green) samples. ***(K)*** Number of alternative splice sites for each gene (in Gencode mouse v14) plotted against gene length. ***(L)*** Distribution of gene lengths for low OFF noise genes (Figure 4B red dashed region) and high OFF noise genes (Figure 4B blue dashed region). Red lines are medians and whiskers indicate 1.5 IQR. (***:p<1e-100, Student's t-test.) ***(M)*** Similar to Figure 4 Supplement 1D, mean Pearson's correlation coefficients between genes within long and short gene groups relative to mean and S.D. (green solid lines) and 99% confidence interval (green dashed lines) calculated from randomly selected groups of genes.

**Figure 7-Supplement 1.**
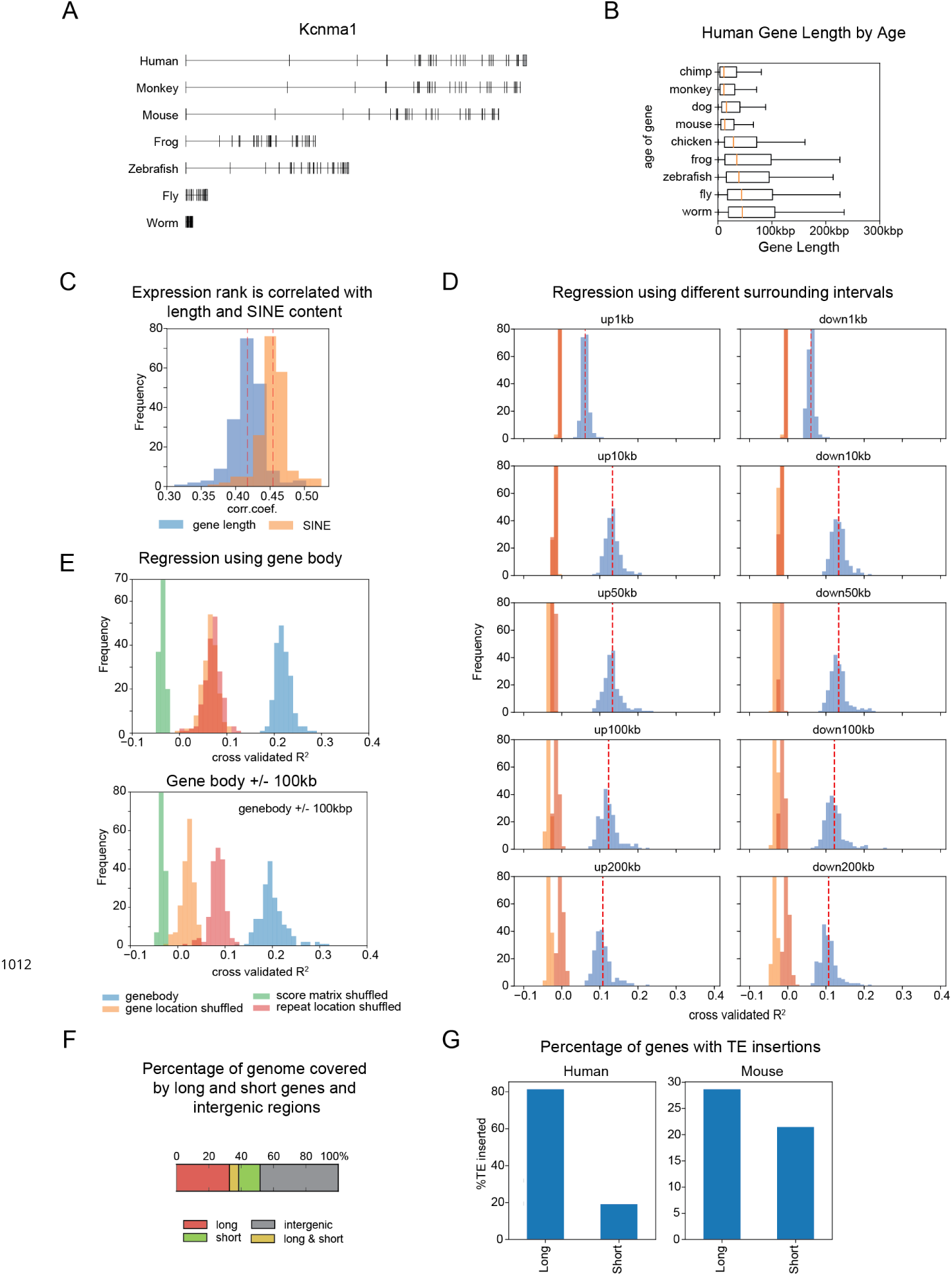
Supplementary to Figure 7. TE insertions elongate genes and contain information about gene expression. **(A)** Example of gene length differences between species for ***Kcnma1*** (a calcium-activated potassium channel, also called *slopoke,* in *Drosophila)*. ***(B)*** Estimated evolutionary age of human genes correlates with their length. The length distribution of human genes is plotted as a function of age, estimated from their most distant homologs. Genes common to all vertebrates (or to all listed genomes) are longer than genes common only to mammals (mouse) or common only to primates (monkey). ***(C)*** Correlation between gene expression rank and gene length (blue) and SINE repeat score (orange) calculated for all cell types. Because of their abundance, SINE repeat scores are correlated with gene length. ***(D)*** Similar to Figure 7E but using repeat scores calculated from different sized intervals surrounding each gene (not including the gene body). Average *R*^2^ is maximal near 10kb for both upstream and downstream intervals. Shuffling conditions are colored as in Figure 7E. ***(E)*** Similar to Figure 7E but for repeat scores calculated from gene body only (upper panel) or gene body+/-100kb (lower panel). ***(F)*** Fraction of genome spanned by long genes (orange) is greater than that spanned by short genes (green), despite being fewer in number. Some genomic regions contain overlapping long and short genes (yellow). ***(G)*** Percentage of inserted sequences calculated in Figure 7A (Human vs. Chimp and Mouse vs. Rat), that overlap TEs within long (***≥*** 100kbp) or short (<100kbp) genes.

## Footnotes

1 A Schur-convex function is a function 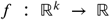 which satisfies ***f (x) ≥ f (y)*** for all ***x,y*** where ***x*** majorizes *y*. For 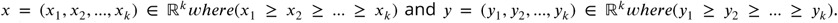. x majorizes *y* when 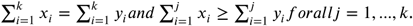. When x majorizes y, it follows *x*_*i*_ ≥ *y*_*i*_ f or all i, so it is easy to see *h(x)* **≥** *h(y)* and *d(x)* **≥** *d(y)*.

